# Noninvasive and Objective Near Real-Time Detection of Pain Changes During Tonic Fluctuating Noxious Heat Stimulation

**DOI:** 10.64898/2026.01.26.701710

**Authors:** Leonard Visser, Christian Büchel

## Abstract

Chronic pain involves natural intensity fluctuations that patients cannot control, contributing to learned helplessness and functional impairment. Detecting spontaneous pain decreases could enable precisely timed interventions that enhance perceived control. However, existing research on objective pain assessment has focused primarily on estimating static intensity from short, predictable stimuli rather than detecting moment-to-moment changes in ongoing pain. This study investigated whether pain decreases can be detected objectively using easily obtainable physiological signals during fluctuating pain. We recorded multiple physiological signals (8-channel EEG, electrodermal activity, heart rate, pupil diameter, facial expressions) from 42 healthy participants (*M*_age_ = 26.2 years, *SD* = 5.1) during calibrated tonic noxious heat stimulation on the left forearm with unpredictable intensity fluctuations (0–70 on visual analogue scale; twelve 3-minute trials). Temperature changes lasted 5–20 seconds. Using minimal preprocessing suitable for real-time applications, we trained deep learning models to classify pain decreases versus non-decreases from brief temporal windows, evaluated on a held-out test set (9 participants). Combining electrodermal activity, heart rate, and pupil diameter yielded optimal classification performance using a transformer-based architecture (AUROC = 0.854, accuracy = 76.8%). Electrodermal activity emerged as the most informative single predictor. Continuous stream analysis demonstrated median detection latency of 5.75 seconds with 70.4% sensitivity, reducible to 4.25 seconds at the cost of increased false positives. Results indicate that electrodermal activity and heart rate enable straightforward practical deployment, while highly variable signals such as EEG and facial expressions require personalized fine-tuned models. These findings establish a basis for closed-loop interventions targeting spontaneous pain changes.

**Summary:** Deep learning models detect pain decreases using electrodermal activity, heart rate, and pupil diameter (AUROC=0.854, 5.75s latency), facilitating precisely timed interventions for fluctuating tonic pain.

## Introduction

Chronic pain impairs patients’ sense of control over their symptoms, contributing to functional disability and fostering learned helplessness [61,74]. Since pain self-efficacy predicts better clinical outcomes, restoring perceived control represents a meaningful therapeutic target [2,3,41,64]. However, patients with chronic pain characteristically report little to no control over their symptoms.

A recently proposed approach [16] suggests that patients may be able to regain at least some degree of perceived control over their pain through a so-called “illusion of control” [54]. The core idea is that if a spontaneous reduction in naturally fluctuating pain is detected and patients are then allowed to treat themselves while the pain diminishes, this may be perceived as a form of causal intervention. Given that the pain is already decreasing, a precisely timed intervention, for example pressing a blinking button to trigger transcutaneous electrical nerve stimulation (TENS), could be sufficient to reinforce this perception [29]. Over time, two therapeutic mechanisms might be engaged: through conditioning, the initiation of the treatment may gradually take on analgesic properties of its own [17,21,24,46], and through the repeated experience of acting and observing relief, patients may perceive greater control over their pain, strengthening their self-efficacy [2,9,10,43]. As a first step, however, this approach requires an instrument that can objectively and noninvasively track ongoing pain levels and reliably detect pain decreases in real time.

Previous research on objective pain assessment has largely focused on brief, discrete, and static noxious stimuli with clearly defined onsets and offsets, aiming to estimate absolute pain intensity from concurrently recorded physiological signals [26,27,32,35,38,44,57,59,60,79,81,84]. Recent advances have introduced state-of-the-art trans-former-based deep learning architectures for this task [25,33], including a cross-modal transformer model combining electrodermal activity (EDA) and electrocardiogram signals [26]. However, these approaches do not adequately capture the moment-to-moment fluctuations characteristic of ongoing, tonic pain. This is particularly important because many chronic pain conditions exhibit natural fluctuations that lie outside patients’ control [7,31,65,66].

In this study, we investigated whether pain dynamics in healthy participants can be assessed objectively using various easily obtainable physiological measures. Our exploratory multi-modal analysis focused on EDA, heart rate, pupil diameter, electroencephalography (EEG), and facial expressions during controlled exposure to tonic, unpredictably fluctuating, noxious heat stimuli. Through systematic evaluation of different feature sets and deep learning model architectures, we identified optimal approaches for detecting pain dynamics, particularly pain decreases. Finally, we validated the performance of our model through a continuous data stream analysis, in which we applied a sliding window technique to approximate real-time application conditions. Taken together, our findings provide proof of concept for objective detection of pain decreases in fluctuating, experimental tonic pain and establish a foundation for subsequent validation in chronic pain populations.

## Methods

### Participants

#### Eligibility Criteria

Participants were eligible if they were healthy, right-handed, between 18 and 40 years of age, and had a body mass index (BMI) between 18 and 30 kg/m². The age and BMI ranges were restricted to reduce physiological heterogeneity unrelated to the experimental manipulation. All participants possessed full legal capacity to provide informed consent and demonstrated adequate German language proficiency for following complex instructions.

Exclusion criteria encompassed chronic pain conditions (Patient Health Questionnaire (PHQ-15) [50], cutoff 10), elevated depressive symptoms (Beck Depression Inventory II (BDI-II) [39], cutoff 14), and other psychiatric, neurological, or cardiovascular conditions. Additional exclusions targeted factors with direct methodological implications for multimodal psychophysiological recording: dermatological abnormalities on the left inner forearm (scarring, tattoos, skin conditions; affecting sensor adhesion and conductance), hair characteristics incompatible with full-scalp EEG electrode placement (natural baldness, completely shaved head, dreadlocks, braided hairstyles), recent Botox facial treatments (within 6 months; attenuating facial muscle movements detectable via video-based expression analysis), facial tattoos or piercings (interfering with facial expression analysis), a need for glasses or hard contact lenses (precluding reliable pupillometry), respiratory conditions (cold symptoms, pollen allergies; potentially confounding autonomic measures), pregnancy or breastfeeding, and regular medication use (except contraceptives). The age range (18–40 years) and BMI range (18–30 kg/m²) were selected to minimize physiological variability attributable to age-related comorbidities and adipose tissue effects on surface electrode recordings, respectively.

#### Participant Characteristics

We enrolled 50 healthy right-handed participants with normal or corrected-to-normal vision (via soft contact lenses). Participants received monetary compensation for their time. Prior to participation, participants reported abstaining from pain medication and cannabis or similar substances for 24 hours, from alcohol for 12 hours, and from caffeine for 2 hours.

#### Participant Flow and Final Sample

Five participants were excluded from data analysis (due to inattentiveness, pain threshold issues, or BMI criteria). From the remaining 45 participants, 69 of a total of 540 trials were excluded for technical reasons (EDA sensor issues, thermode malfunctions, EEG recording problems) or protocol violations, as detailed in Figure 1. The final dataset thus comprised 471 valid trials from 42 participants (23 women; age range from 18 to 39 years; mean 26.2).

**Figure 1:**
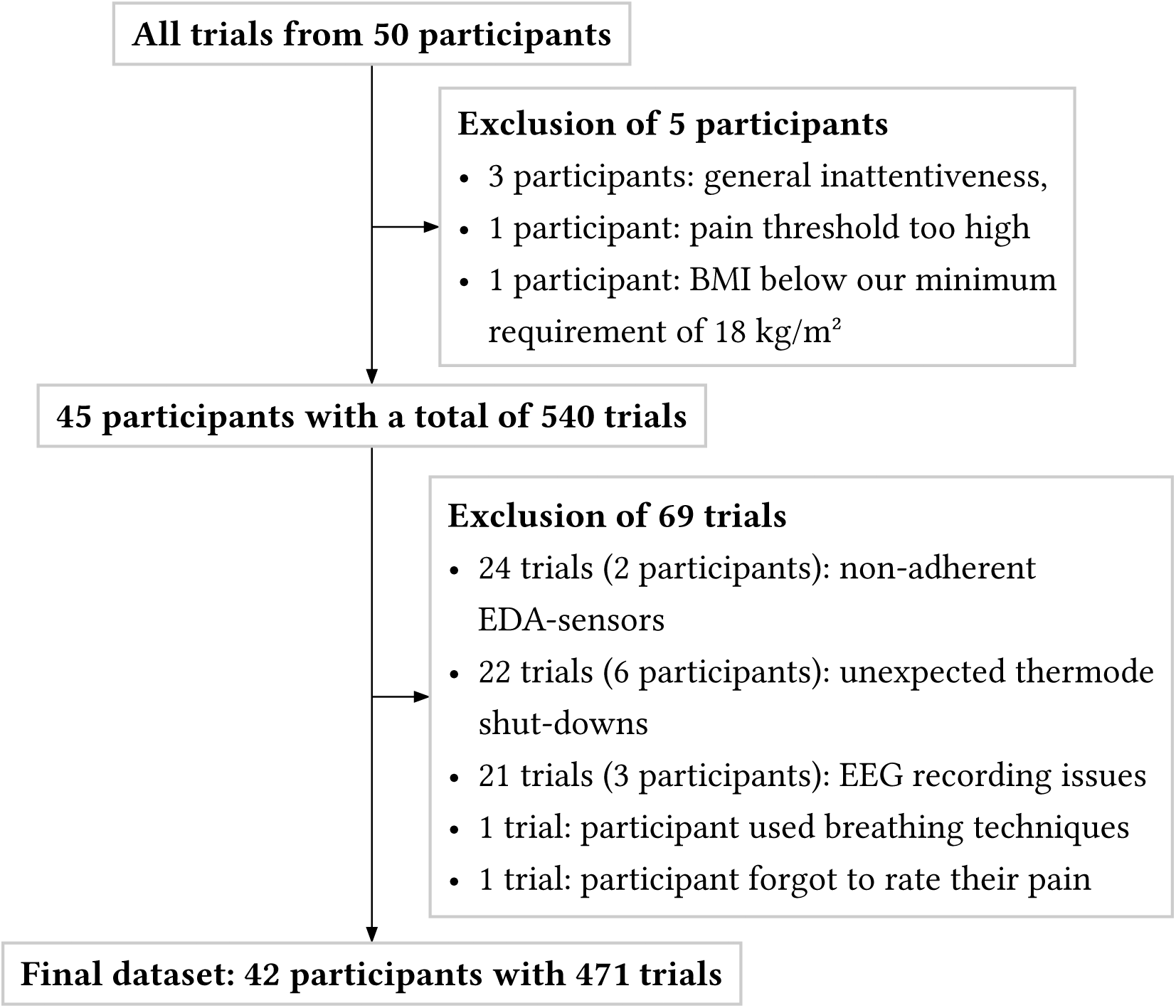
Flow diagram of participants and experimental trials. Exclusion criteria were applied in two stages: first at the participant level for eligibility requirements, then at the trial level for data quality and protocol adherence. BMI = body mass index, EDA = electrodermal activity, EEG = electroencephalography.

The Ethics Committee of the Medical Chamber Hamburg approved the study (procedure number 2023-101089-BO-ff), and all participants provided written informed consent. Data collection took place from June to December 2024 at the Department of Systems Neuroscience, University Medical Center Hamburg-Eppendorf.

### Noxious Heat Stimulus

To evoke fluctuations in tonic pain, we used sinusoidal functions to create controlled temperature curves of 180 s length with unpredictable patterns. Each stimulus function consisted of 5 ascending and 5 descending segments with different amplitudes and periods (5 to 20 s). Individual segments were generated using half-cycle cosine waveforms following the equation

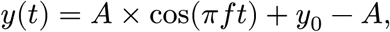

where *A* is amplitude, *f* is frequency, and *y*_0_ is the *y*-intercept. These segments were joined in an alternating up-and-down pattern (switching between positive and negative values) to form a smooth, continuous temperature curve of 145 s. To further prevent anticipation effects, three temperature plateaus were added: two plateaus of 15 s length within ascending segments and one plateau of 5 s length at the point of a local minimum, yielding a total length of 180 s (see Suppl. Figure S1 for details). All heat stimuli included three uniform major decreasing intervals to facilitate our primary research focus on detecting pain decreases (see Figure 2 for an example).

**Figure 2:**
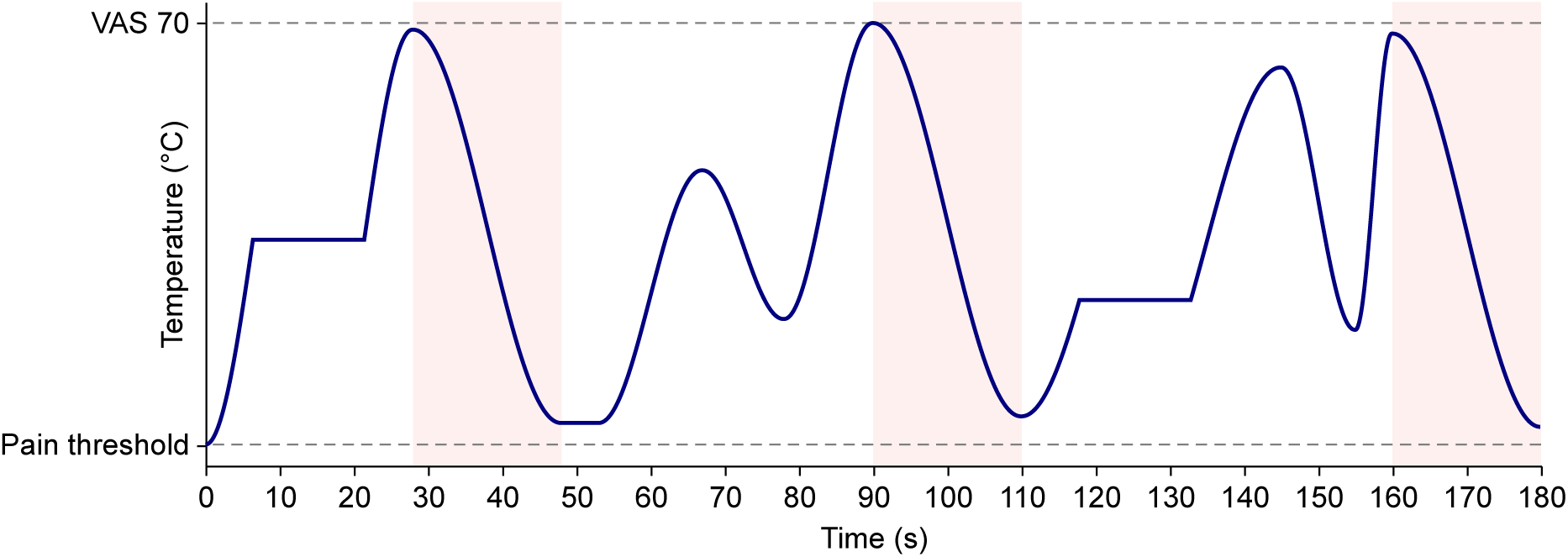
Example of a temperature curve for one trial. Each participant completed a set of 12 three-minute trials with distinct temperature curves. We scaled each curve to the participant’s calibrated range from pain threshold to the temperature corresponding to a rating of 70 on the visual analogue scale (VAS). Red regions mark the three standardized major decreasing intervals that supplied decrease-class samples. Increasing and plateau intervals supplied non-decrease samples, as detailed in Suppl. Figure S1.

Each participant received 12 unique 180 s stimuli (see Suppl. Figure S2 for a visualization of all stimulus curves), with pain intensities fluctuating between pain threshold and strong pain (70 out of 100 on a visual analogue scale (VAS)). While the stimulus patterns remained consistent across participants, we individually calibrated temperatures (see below for details) to each participant’s pain threshold and VAS 70. The presentation order of the stimuli was randomized.

### Procedure

We collected multiple psychophysiological measures along with online pain ratings while applying the heat stimuli. Psychophysiological measures included EDA, pupillometry, photoplethysmography for heart rate (HR), EEG, and facial expressions through video recordings. The experiment utilized a within-subject design and lasted approximately 2.5 hours per participant. All participants were tested individually by the same experimenter in a controlled laboratory environment with standardized illumination. Participants sat in a comfortable chair with a computer monitor at eye level, positioned approximately 65 cm away. The experimenter provided standardized instructions with limited interaction. The experiment consisted of five phases: (1) pre-experiment questionnaires, (2) experiment set-up, (3) pain calibration, (4) pain measurement, and (5) post-experiment questionnaires. These phases are outlined in the following sections.

#### Pre-Experiment Questionnaires

Following a brief introduction and informed consent procedure, participants completed psychological screening questionnaires: a general questionnaire assessing socioeconomic status, the BDI-II, and the PHQ-15. To enable comparison of mood states before and after the heat stimulation, participants also completed the Positive and Negative Affect Schedule (PANAS) [14].

#### Experiment Set-up

After completing the questionnaires, participants washed their hands without soap to optimize skin conductance electrode attachment. We then applied measurement equipment in the following sequence:

1. An 8-channel EEG system (Neuroelectrics Enobio 8) with a 500 Hz sampling rate was placed with electrodes at positions F3, F4, C3, CZ, C4, P3, P4, and OZ according to the international 10-20 system. We used Signal Gel electrode gel, with reference electrodes (DRL and CMS; Cardinal Health Kendall H124SG) placed on the previously cleaned left mastoid.
2. Electrodermal activity and photoplethysmography (PPG) for heart rate were recorded using a Shimmer3 GSR+ device with a 100 Hz sampling rate. EDA electrodes (3M Red Dot 2239) were attached to the upper and lower portions of the left palm below the little finger, and the PPG sensor was attached to the left earlobe.
3. Pupil diameter was measured using an eye tracker (Smart Eye AI-X, 60 Hz) mounted on the lower part of the monitor.
4. Facial expressions were recorded using a high-definition camera (Logitech C920s PRO, 1024 × 768 pixel resolution).

Signal quality was visually validated for all physiological measures, and eye tracker calibration was performed before proceeding to the pain calibration. Recording of all physiological measures commenced at this point and continued throughout the remainder of the experiment.

The noxious heat stimulation was applied to six distinct skin patches on the participant’s left forearm, organized in two parallel columns of three positions each according to a standardized template. To minimize habituation and sensitization effects, stimulation alternated between patches, maximizing the distance between consecutive stimulations. Each patch received two stimulations throughout the experiment. The stimulation order was counterbalanced across participants (see Suppl. Figure S3 for details on the counterbalancing scheme). All stimuli were applied using a Pathways thermode (Medoc Ltd., Israel) with an ATS thermode (30 × 30 mm²).

#### Pain Calibration

We calibrated two individual stimulation temperatures: pain threshold and VAS 70. The heat stimuli lasted 9 s at the plateau with a 10°C/s rise rate from a 32°C baseline and randomized inter-trial intervals of 12–13 s. The protocol began with warm-up stimuli (39°C, 40°C) and pre-exposure temperatures (42°C, 43°C, 44.5°C) to assess initial pain sensation. The following pain threshold calibration employed recursive Bayesian updating with a Gaussian prior (*μ* = 43°C if pre-exposure pain reported or else *μ* = 44.5°C, *σ* = 1.75°C). Participants provided binary above/below pain threshold feedback for 6 trials, with progressively decreasing likelihood variance to reflect increasing estimation confidence. The threshold was set as the posterior distribution maximum. VAS 70 calibration used the same method with the prior centered at the pain threshold plus 2.5°C (*σ* = 0.65°C). To ensure participant safety, we set the maximum temperature to 48°C. We further enforced a minimum 1.5°C difference between threshold and VAS 70 to maintain a sufficient temperature range, adjusting the calibrated temperatures if necessary while preserving safety limits. The approximately 5-minute-long calibration yielded average temperatures of 44.5°C (*SD* = 1.5°C) for pain threshold and 46.8°C (*SD* = 0.8°C) for VAS 70 before any adjustments.

#### Pain Measurement

Participants received 12 unique noxious heat stimuli, each lasting 3 minutes. The temperature fluctuated between the individually calibrated pain threshold and VAS 70 values. Throughout each trial, participants continuously rated their pain intensity by moving the computer mouse with their right hand on a digital, equiluminant VAS ranging from no pain (“No pain”) to strong pain (70 VAS; “Very strong pain”). Participants were instructed to keep their left arm with the thermal stimulation and EDA device still, to fully concentrate on their pain experience, and to avoid distraction. Starting from the baseline temperature of 32°C, we increased the temperature at a rate of 10°C/s until reaching each participant’s individual pain threshold. After reaching the pain threshold, we introduced a pause of 1.5 s to display the interactive VAS rating scale, without recording ratings. Following this pause, we initiated the tonic, fluctuating noxious heat stimulus while simultaneously starting the recording of online pain ratings. Pain ratings were sampled at a frequency of 10 Hz. To increase motivation, participants were rewarded with 0.50 € if the correlation between their rating and the applied temperature was at least 0.7 in a single trial. This threshold was intentionally set to be easily achievable so as not to add unnecessary cognitive load to the task. Out of 471 trials, 315 trials were rewarded. Between trials, we moved the thermode to the next skin patch. Throughout the pain measurement, all physiological signals were continuously monitored by the experimenter to ensure high data quality. The measurement lasted approximately 45 minutes.

#### Post-Experiment Questionnaires

Following the pain measurement, participants completed the PANAS a second time and additional questionnaires: the Pain Catastrophizing Scale (PCS) [78] for pain catastrophizing, the Pain Vigilance and Awareness Questionnaire (PVAQ) [52] for pain attention and vigilance, the State-Trait Anxiety Inventory-Trait version (STAI-T-10) [75] for trait anxiety, and the Mindful Attention Awareness Scale (MAAS) [15] for dispositional mindfulness. Individual questionnaire results are provided in Suppl. Table S1 to Table S4. Participation in the experiment was compensated with 37–43 €, based on the individual rating performance in the pain measurement task.

#### Data Acquisition

The experiment was programmed using the Python library Expyriment (version 0.11.0.dev1) [48], which managed stimulus presentation and online pain ratings. Synchronization and recording of all behavioral and psychophysiological data were coordinated by the iMotions software suite (version 10.0) [40] in conjunction with the Neuroelectrics Instrument Controller software (version 2.1.3.6) for EEG data. For further analyses and efficient data processing, we stored the CSV exports from iMotions in the SQL database system DuckDB (version 1.2.2) [73]. All original participant IDs were randomly reassigned to new, anonymized IDs.

### Preprocessing

For the classification task, our preprocessing pipeline was intentionally kept minimal to support potential real-time deployment. Transformations were restricted to causal transformations, i.e., methods that use only current and past data points. This constraint excluded more sophisticated preprocessing steps, such as EDA deconvolution. For EEG data, we did not apply eye-blink artifact correction or similar procedures, as raw EEG data has been shown to deliver robust results in combination with deep learning analysis [23]. For an additional exploratory data analysis, we utilized non-causal transformations to better preserve the temporal properties of the original signals.

A direct comparison between the two approaches was precluded by the inherently dynamic nature of the task, as non-causal operations incorporate future information and thus would introduce data leakage in the classification task. Some offline preprocessing approaches (e.g. certain ICA artifact removal techniques), however, may still be informative as complementary robustness checks or for physiological source attribution in future work.

In a final preprocessing step, we decimated all physiological signals except EEG data to a sampling rate of 10 Hz and merged them into a single feature matrix with equidistant time steps. EEG data were processed separately at a sampling rate of 250 Hz.

#### Electrodermal Activity

For the classification task, we used raw EDA data obtained directly from the Shimmer3 device. In line with the causal real-time objective, abrupt skin conductance changes potentially related to movement were not explicitly removed or smoothed. For the exploratory data analysis, we separated tonic (slow changes) and phasic (fast changes) components using a 5th-order Butterworth low-pass filter with a cutoff frequency of 0.05 Hz, implemented using NeuroKit2 (version 0.2.10) [62].

#### Pupillometry

Blink periods were identified using the eye tracker’s native detection algorithm. The pupillometric data underwent additional preprocessing steps to further improve signal quality. First, physiologically oriented thresholds were applied to remove pupil size measurements falling outside a 2–8 mm range. For the causal preprocessing, identified blink periods were filled using forward propagation. For the exploratory data analysis, identified blink periods were extended by 120 ms on both sides to exclude measurement artifacts that frequently occur at the boundaries of blink events [49]. A median filter with a 1 s local window was then applied to remove partially detected blinks that remained in the signal. For the classification task, the median filter was implemented with forward padding, ensuring each output sample depended only on current and previous samples. For the exploratory data analysis, the filter was symmetric. Finally, the right and left pupil measurements were averaged into a single dilation measure.

#### Heart Rate

Stepwise heart rate data was acquired directly from the Shimmer3 device. To ensure physiological plausibility, we imposed an upper threshold of 120 beats per minute. This threshold was used as a physiological plausibility cutoff for resting seated participants. For causal preprocessing, missing values were identified and filled using forward filling. For exploratory data analysis, we utilized linear interpolation to fill the gaps in the time series and applied a second-order Butterworth filter with a highcut of 0.8 Hz to mitigate abrupt transitions in the signal.

#### Facial Expressions

Over 30 different features were extracted using the Affectiva iMotions Facial Expression software AFFDEX 2.0 [11], which has been demonstrated to reliably identify facial expressions with accuracy comparable to traditional electromyography (EMG) measurements [51]. Based on previous research on pain-relevant facial expressions [53,72], we focused our analysis on five key features: brow furrow, cheek raise, mouth open, nose wrinkle, and upper lip raise. These features were represented as probability values between 0 and 1, indicating the likelihood of the expression being present. The raw data were preprocessed causally using an exponential moving average smoothing with an alpha of 0.05; for non-causal preprocessing we used a second-order Butterworth filter with a highcut of 0.2 Hz.

#### Electroencephalography

We preprocessed EEG signals causally through a three-stage pipeline using SciPy (version 1.15.2) [83]. All filters were implemented as forward-only IIR filters. Data were first decimated from 500 Hz to 250 Hz utilizing a 6th-order Butterworth anti-aliasing filter. We then applied a 4th-order Butterworth high-pass filter with a 0.5 Hz cutoff to remove slow drifts. Finally, second-order notch filters (quality factor of 30) at 50 and 100 Hz eliminated power line interference and its first harmonic. A detailed exploratory analysis of the EEG data was beyond the scope of the current study.

### Exploratory Data Analysis

#### Psychophysiological Signals

To examine relationships between physiological measures, we created grand-averaged, min-max scaled time series with a bin size of 0.1 s and 95% confidence intervals for each measure (pain ratings, tonic and phasic EDA, pupil diameter, heart rate, facial expressions) by averaging across all participants for each temperature curve. We calculated Pearson correlation coefficients between these grand-averaged time series to create two correlation matrices, one for the psychophysiological signals and one for the facial expressions. Additionally, we performed cross-correlation analyses to examine time-lagged relationships for the psychophysiological measures with the stimulation temperatures. To evaluate the reliability of these patterns on participant level, we calculated individual-level mean correlations across the 12 stimulus functions, then computed summary statistics (overall mean, standard deviation, minimum, and maximum) of these participant-level means across the entire sample. Correlations were averaged directly without Fisher z-transformation.

For all above-mentioned correlations, we excluded the first 20 s of each trial to ensure physiological equilibrium. This exclusion was necessary because the steep initial temperature ramp-up from baseline to pain threshold (>10°C increase in under 3 s) created artifacts unrelated to subsequent stimulation, particularly affecting low-pass filtered tonic components. The 20 s exclusion period was empirically determined from grand-averaged data, which showed stable responses after this timepoint, enabling reliable assessment of pain-related physiological dynamics.

#### Pain Ratings

We conducted exploratory analyses focused specifically on pain ratings. First, to assess potential learning effects, we computed Pearson correlations between individual pain ratings and temperature curves for each 180 s-long trial and examined whether these correlations changed systematically over the course of the experiment. Second, to evaluate whether participants perceived the full calibrated range of stimulus intensity without habituation effects, we extracted the maximum and minimum pain ratings within each major decreasing temperature interval across trials. Third, to specifically characterize the coupling between pain ratings and temperature during major decreases, we computed grand-averaged pain ratings.

### Deep Learning Analysis

#### Sample Creation

We created 2826 samples of 7 s length for the binary classification task discriminating between pain decreases and non-decreases. For the non-decrease category, we utilized two types of temperature patterns: increases and plateaus. Temperature increases consisted of the first 7 s of intervals with strictly increasing temperatures that contained no plateaus in them (see Figure 2). We excluded the first increasing temperature interval from each trial. Temperature plateaus were based on the 15 s periods of constant temperature stimulation, from which we extracted one 7 s sample starting at a 5 s offset. For pain decreases, we shifted the samples by 1 s from the start of a major decreasing interval to account for a latency in pain perception following the temperature reduction after the temperature peak. This window was chosen to capture the early phase of the decrease as timing is important for the proposed intervention. The same 7 s window length was applied across modalities to provide a shared temporal context for multimodal classification. All samples were constructed as non-overlapping temporal windows.

Decreases followed a uniform temperature pattern due to the standardized curves of major decreases in the stimulus function. In contrast, pain increases exhibited more variability in amplitude and slope, and plateaus had different levels. Since more samples were available for non-decreases, we balanced the labels by downsampling the majority class to match the minority class (decreases). This ensured the models were trained on a balanced dataset, which is crucial for effective classification performance [18]. The final dataset comprised 2826 samples, with 1413 samples per class.

#### Data Splitting Strategy

To prevent participant-dependent data leakage, we implemented a group-based data splitting strategy. Participants were assigned exclusively to training, validation, or test sets in a 60:20:20 ratio. This resulted in 1615 samples (24 participants) for training, 626 samples (9 participants) for validation, and 585 samples (9 participants) for the test set. We standardized each feature independently via z-score normalization using parameters obtained from training samples only. For final model training, standardization parameters were computed from the combined training and validation sets.

#### Feature Sets

We systematically evaluated 11 physiological feature sets, organized hierarchically to reflect increasing feature set complexity. We began with individual modalities: (1) EDA, (2) HR, and (3) pupil diameter. We then combined peripheral autonomic signals into composite sets: (4) EDA + HR, (5) EDA + pupil diameter, and (6) EDA + HR + pupil diameter. Next, we introduced behavioral measures by adding (7) facial expressions alone and (8) facial expressions combined with all peripheral signals. Finally, we incorporated the central nervous system measure by evaluating (9) EEG alone, followed by (10) EEG + EDA and (11) EEG combined with all remaining modalities (facial expressions + EDA + HR + pupil diameter). A combination of EEG and HR was not evaluated, as both modalities individually performed just above chance level under the real-time preprocessing constraints described above.

#### Model Architectures

For non-EEG signals (feature sets 1–8), we evaluated five distinct neural network architectures representing different modeling approaches: a multilayer perceptron (MLP) as a baseline reference [56], LightTS (MLP-based with different downsampling strategies) [88], TimesNet (CNN-based with period transformation) [85], PatchTST (transformer with patch-based tokenization) [69], and iTransformer (transformer with variate-based tokenization) [58].

For EEG data (feature sets 9–11), we employed a different set of architectures, since not all previously mentioned models scaled effectively to time series with higher temporal resolution. We evaluated an MLP, LightTS, iTransformer, and EEGNet, a convolutional neural network architecture specifically designed for EEG signal analysis [55].

We additionally implemented feature-level fusion ensemble models [8], combining separate PatchTST models for facial expressions and peripheral physiological signals (set 8), EEGNet for EEG with PatchTST for physiological signals (set 10), and EEGNet for EEG with PatchTST models for both facial expressions and physiological signals (set 11). Detailed architectural descriptions, including modality-specific sampling rates, are provided in Suppl. Material C.

#### Model Training and Evaluation

The factorial design of 11 feature sets and their respective model architectures resulted in 53 distinct model-feature combinations. Following the standard train-validation-test split method, we conducted hyperparameter optimization for each combination using Bayesian-inspired search via the Optuna library [1] to maximize validation accuracy. After identifying the optimal configuration for each model architecture, we retrained the best-performing model for each feature set on the combined training and validation set before conducting a final evaluation on the previously unseen test set. By doing so, we maximized the training data available to each final model while preventing data leakage. This process yielded 11 final models, one optimal configuration for each feature set.

We evaluated overall test set performance using accuracy, area under the receiver operating characteristic curve (AUROC), precision, recall, F₁-score, and Matthews correlation coefficient (MCC), with pain decreases defined as positive cases. We generated receiver operating characteristic (ROC) curves to visualize discriminative performance across feature sets. Additionally, we evaluated individual-level test accuracy for each participant across all feature sets to assess prediction consistency. Full details on hyperparameter search spaces, training procedures, and metric definitions are provided in Suppl. Material D.

#### Seed Stability Analysis

To assess the stability of results across different participant-based group splits, we performed a seed stability analysis. Using the best-performing model configuration, we evaluated performance across multiple random train-test splits (*n* = 200) produced by varying the random seed. For each split, we trained the model on the combined training and validation sets, then evaluated on the test set. We visualized the distribution of the test accuracies across all seeds using histograms for the feature sets EDA + HR and EDA + HR + pupil diameter.

#### Application to Continuous Data Streams

We applied the best-performing models to continuous data streams to detect pain decreases in fluctuating pain. Unlike the training set, where we used non-overlapping samples specifically selected from clearly defined temperature intervals, we created a new set of samples based on the full 180 s trials from participants in the test set. This approach simulates real-world conditions, where the model must continuously evaluate pain dynamics on a moment-to-moment basis. Using a sliding window approach with 7 s windows and a 250 ms step size between samples, we created a total of 693 samples for each trial and participant in the test set. Samples were standardized with values obtained from the combined training and validation set. No additional training was performed.

From the model’s output, we converted the raw logits into probability values using a sigmoid function. We visualized them as confidence values for detecting decreases across time for each participant in a heatmap. Only decreases were visualized for visual clarity. For the best-performing model, the Supplementary Materials additionally provide prediction heatmaps for non-decreases alone and for both classes combined. We selected conservative classification thresholds to reduce false positives, as high precision would be crucial for ensuring that interventions are triggered primarily during genuine pain decreases. For better interpretability, we also visualized the mean pain ratings of the test set participants within 95% confidence intervals.

To determine the latency of the classifier, we calculated the median time and median absolute deviation (MAD) of the first detection of pain decreases in major temperature decreases. Detections occurring within 1 s after the start of a major temperature decrease were removed for physiological plausibility.

## Results

### Exploratory Data Analysis

#### Psychophysiological Signals

##### Grand-Average Level

Correlation analysis of grand-averaged data revealed systematic relationships between temperature curves, pain ratings, and physiological measures (for visualized grand-averages, see Figure 3A; for a correlation matrix, see Figure 3B). Temperature and pain ratings showed the strongest overall correlation (*r* = .89). Among physiological measures, pupil diameter demonstrated the highest correlation with temperature (*r* = .61), followed by heart rate (*r* = .42), tonic EDA (*r* = .39), and phasic EDA (*r* = .19). A visualization of the grand-averaged signals over all 12 temperature curves is provided in Suppl. Figure S4.

**Figure 3:**
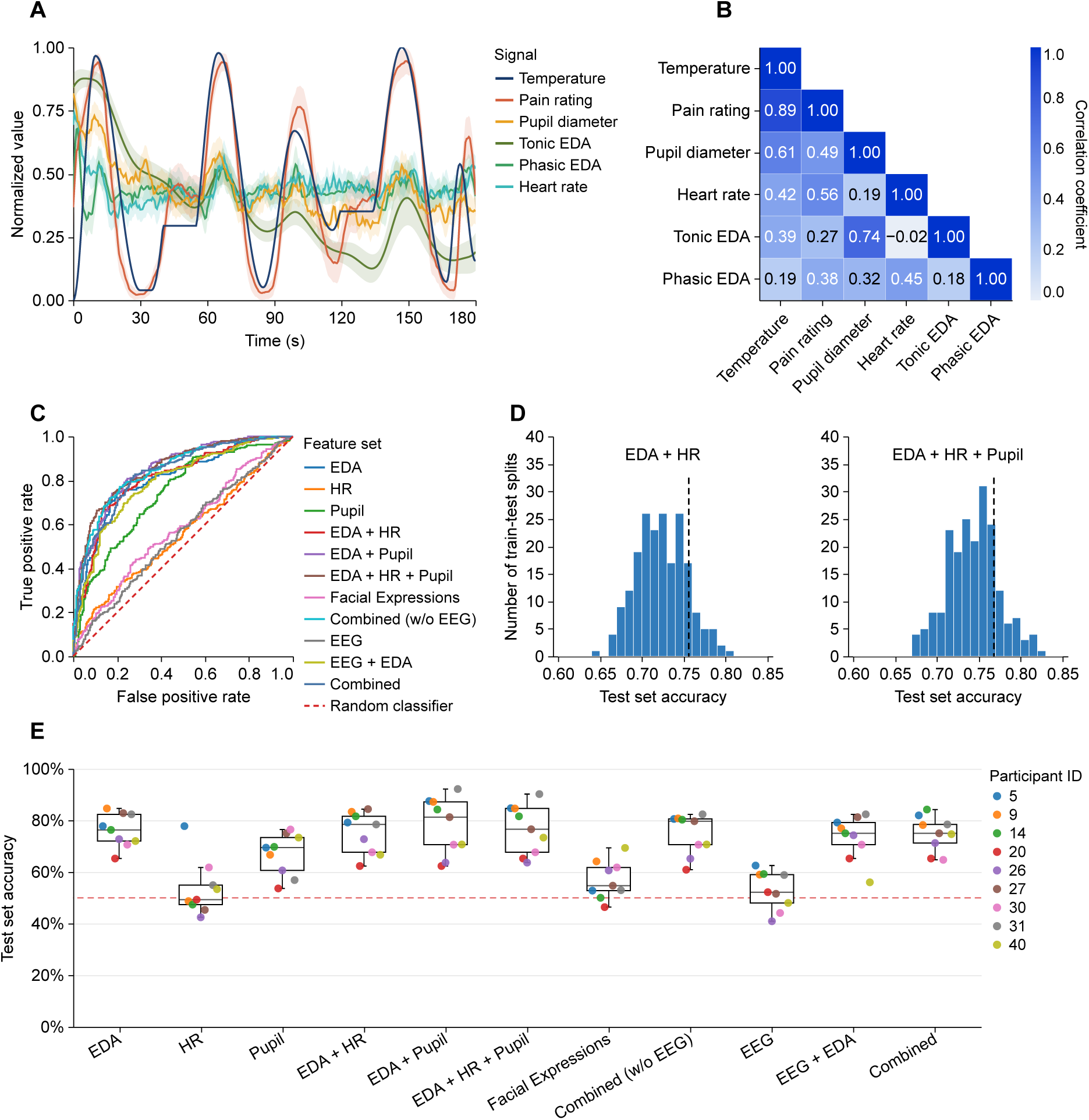
Exploratory data analysis, model performance, and individual differences. **A** Grand-averaged temperature, continuous pain ratings, and physiological signals during one example 180-second stimulus curve. We normalized each signal to the range 0–1 for visualization; shaded bands show 95% confidence intervals across participants. **B** Correlation matrix of grand-averaged temperature curves, pain ratings, and physiological measures. The heatmap shows Pearson correlation coefficients between signals, with darker blue indicating stronger positive correlations. **C** Receiver operating characteristic (ROC) curves for models classifying 7-second physiological windows from major decreasing temperature intervals versus windows from increasing and plateau temperature intervals using physiological signals (without pain ratings). The dashed diagonal marks chance-level discrimination. **D** Test-accuracy distributions across 200 random participant-based data splits for the EDA + HR and EDA + HR + pupil-diameter feature sets. Dashed vertical lines mark the accuracies obtained with the primary data split. **E** Test accuracy across the nine held-out participants for each of the eleven feature sets. Points represent participants, boxes show the median and interquartile range, and the red dashed line marks chance-level (50%) accuracy. EDA = electrodermal activity; HR = heart rate; “Facial expressions” comprise video-based detection of brow furrow, cheek raise, mouth open, nose wrinkle, and upper lip raise; EEG = electroencephalography; “Combined (w/o EEG)” comprises EDA, HR, pupil diameter, and facial expressions; “Combined” comprises those measures along with EEG.

Regarding facial expressions, cheek raise showed a moderate correlation with temperature (*r* = .60), followed by mouth open (*r* = .39) and upper lip raise (*r* = .27). Nose wrinkle (*r* = .09) and brow furrow (*r* = .05) were only minimally correlated with temperature. All grand-averaged facial expression activations remained largely near baseline (normalized value of 0) throughout the full stimulation (for visualized grand-averages, see Suppl. Figure S5; for a correlation matrix, see Suppl. Figure S6).

Cross-correlation analysis revealed minimum lags behind temperature stimulation for pupil responses (*M* = 0.07 s, *SD* = 0.22 s), followed by tonic EDA (*M* = 0.95 s, *SD* = 0.62 s), pain rating (*M* = 1.01 s, *SD* = 0.34 s), heart rate (*M* = 1.76 s, *SD* = 1.36 s), and phasic EDA (*M* = 2.45 s, *SD* = 0.89 s). Additional lags for facial expressions can be found in Suppl. Table S5, and cross-correlation lags with pain ratings as reference are provided in Suppl. Table S6.

#### Individual Participant Level

At the individual participant level, we observed consistent pain-temperature correlations (*M* = 0.77, *SD* = 0.05, ranging from *r* = .68 to *r* = .87) and substantial variability in physiological response correlations. Pupil diameter showed the widest range of correlations with temperature, from *r* = –.18 to *r* = .61 (*M* = 0.26, *SD* = 0.20), followed closely by tonic EDA correlations ranging from *r* = –.09 to *r* = .67 per participant (*M* = 0.21, *SD* = 0.20). Heart rate ranged from *r* = –.15 to *r* = .53 (*M* = 0.12, *SD* = 0.16). Phasic EDA responses were minimal and consistent (*M* = 0.07, *SD* = 0.06). Suppl. Figure S7 provides a graphical representation of these correlations for each participant.

Facial expressions showed high variance across participants, with mean correlations ranging from *r* = –.37 to *r* = .64, mostly distributed around *r* = 0 (see Suppl. Figure S8). Only two participants displayed moderate positive correlations (around *r* = .40) across all five analyzed facial expressions.

#### Pain Ratings

Exploratory analyses of pain ratings revealed consistent psychophysical coupling with the noxious heat stimulation throughout the experiment. Pain-temperature correlations remained stable across all 12 trials, with no notable differences across trials (see Suppl. Figure S9). Within major decreasing temperature intervals, participants consistently utilized the full rating scale from VAS 0 to VAS 70 (see Suppl. Figure S10 and Figure S11).

### Deep Learning Analysis

#### Model Comparison Results

We evaluated 53 model-feature combinations, with performance metrics summarized in Table 1 and corresponding ROC curves shown in Figure 3C. The EDA + HR + Pupil feature set achieved the highest AUROC of 0.854 (accuracy = 0.768) using the PatchTST architecture. The EDA + Pupil combination showed comparable performance with an AUROC of 0.851 (accuracy = 0.779). The third-best performance was achieved with the feature set containing EDA + HR + Pupil + facial expressions, with an AUROC of 0.846 (accuracy = 0.747).

**Table 1:** Model comparison results across different feature sets.

| Feature set | Best model | Acc. | AUROC | Precision | Recall | F<sub>1</sub> | MCC |
| --- | --- | --- | --- | --- | --- | --- | --- |
| EDA | PatchTST | 0.762 | 0.805 | 0.806 | 0.700 | 0.750 | 0.530 |
| HR | MLP | 0.530 | 0.557 | 0.550 | 0.411 | 0.470 | 0.065 |
| Pupil | iTransformer | 0.667 | 0.738 | 0.675 | 0.663 | 0.669 | 0.333 |
| EDA + HR | PatchTST | 0.756 | 0.813 | 0.775 | 0.731 | 0.752 | 0.512 |
| EDA + Pupil | PatchTST | 0.779 | 0.851 | 0.778 | 0.791 | 0.785 | 0.559 |
| EDA + HR + Pupil | PatchTST | 0.768 | 0.854 | 0.762 | 0.788 | 0.775 | 0.535 |
| Facial Expressions | PatchTST | 0.569 | 0.582 | 0.593 | 0.481 | 0.532 | 0.143 |
| Combined (w/o EEG) | PatchTST | 0.747 | 0.846 | 0.726 | 0.805 | 0.764 | 0.496 |
| EEG | iTransformer | 0.535 | 0.555 | 0.538 | 0.593 | 0.564 | 0.069 |
| EEG + EDA | Ensemble | 0.732 | 0.793 | 0.733 | 0.741 | 0.737 | 0.463 |
| Combined | Ensemble | 0.756 | 0.832 | 0.781 | 0.721 | 0.750 | 0.513 |

*Note:* Metrics were computed at a classification threshold of 0.5 with a balanced test set. Major decreasing temperature intervals formed the positive decrease class; increasing and plateau intervals formed the non-decrease class. The best model for each feature set was selected using validation accuracy, retrained on the combined training and validation sets, and evaluated once in the test set. Acc. = accuracy, the proportion of correctly classified samples; AUROC = area under the receiver operating characteristic curve; precision = proportion of predicted decreases that were decrease samples; recall = proportion of decrease samples detected by the model; F<sub>1</sub> = harmonic mean of precision and recall; MCC = Matthews correlation coefficient; EDA = electrodermal activity; HR = heart rate; EEG = electroencephalography; MLP = multi-layer perceptron. Pupil denotes pupil diameter. “Combined without EEG” comprises facial expressions, EDA, HR, and pupil diameter; “Combined” comprises those measures plus EEG.

Individual physiological measures demonstrated varying discriminative capabilities. EDA emerged as the most informative single modality (AUROC = 0.805, accuracy = 0.762), followed by pupil diameter (AUROC = 0.738, accuracy = 0.667). Heart rate (AUROC = 0.557, accuracy = 0.530), facial expressions (AUROC = 0.582, accuracy = 0.569), and EEG (AUROC = 0.555, accuracy = 0.535) demonstrated limited discriminative power at the group level.

The ensemble model combining all physiological measures without EEG achieved an AUROC of 0.846 (accuracy = 0.747), slightly lower than the optimal three-feature combination, while the full ensemble including EEG showed an AUROC of 0.832 (accuracy = 0.756). Across feature sets containing EDA, PatchTST consistently emerged as the best-performing architecture. For univariate EDA data, PatchTST achieved a validation accuracy 7.4 percentage points higher than the next-best architecture (see Suppl. Table S7 for validation set accuracies across all model-feature combinations). The iTransformer architecture proved most effective for EEG analysis. Hyperparameter optimization results can be found in Suppl. Table S9.

#### Seed Stability Results

Across 200 random participant-based train-test splits, the EDA + HR model achieved a mean test accuracy of 0.724 (*SD* = 0.030), while the EDA + HR + Pupil model achieved 0.744 (*SD* = 0.031), as shown in Figure 3D. The test accuracies of our primary models (0.756 and 0.768, respectively) were approximately one standard deviation above the mean across all random splits.

#### Interindividual Differences

Classification accuracy varied substantially across the nine test-set participants (see Fig-ure 3E). The EDA + Pupil feature set demonstrated the highest mean accuracy (*M* = 0.777), along with the widest interquartile range (*IQR* = 0.166) and the highest maximum value (0.922). Five participants achieved accuracies above 0.80 with this feature set, more than with any other feature combination.

Univariate EDA showed relatively consistent performance across participants (*M* = 0.760, *IQR* = 0.104), with seven of nine participants exceeding 0.70 accuracy. Pupillary responses yielded consistent but more moderate accuracy (*M* = 0.669, *IQR* = 0.127). Heart rate features produced the second-lowest mean accuracy (*M* = 0.534, *IQR* = 0.075) with pronounced variability: one participant achieved 0.778 accuracy while the remaining participants ranged between 0.493 and 0.618.

Facial expressions demonstrated limited discriminative capability (*M* = 0.570, *IQR* = 0.090) for most participants, with a maximum accuracy of 0.693. EEG also demonstrated limited discriminative capability (*M* = 0.529, *IQR* = 0.11), with only one participant exceeding 0.60 accuracy.

Notably, for EEG, HR, and facial expressions, classification accuracy sometimes fell below chance level. This was most prominent with EEG data, where one participant obtained an accuracy of 0.409, the lowest accuracy overall. Combining EDA and pupil diameter with EEG and/or facial expressions stabilized performance across participants (Combined: *M* = 0.748, *IQR*

= 0.072), though it did not improve maximum accuracy compared to simpler feature sets. No single feature set uniformly optimized classification across all participants, with individual-specific patterns evident across feature sets. Note that individual participants in the test set had slightly different yet class-balanced sample sizes, which are reported in Suppl. Table S10. Exact values of the boxplot diagram can be found in Suppl. Table S11.

#### Application to Continuous Data Streams

Application of the PatchTST model with EDA + HR + Pupil features to fluctuating temperature curves at a conservative classification threshold of 0.90 produced the prediction heatmap shown in Figure 4. The model achieved a sensitivity of 70.37% for major temperature decreases, detecting 209 out of 297 decreases. The median detection latency was 5.75 s (*MAD* = 1.50 s) within the 20-second windows of temperature decrease. Lowering the threshold to 0.5 decreased median latency to 4.25 s (*MAD* = 1.25 s), illustrating a sensitivity-specificity trade-off with improved detection speed at the cost of increased false positive classifications. The predicted pain decreases demonstrated strong temporal alignment with actual temperature decreases across most participants, identifying temperature decreases that were both shorter in duration and smaller in magnitude than those presented during model training. Temperature plateau regions occasionally resulted in false positive classifications. Participant 40 exhibited almost no positive predictions at the conservative threshold level. Corresponding heatmaps for the other 10 feature sets are shown in Suppl. Figure S12 through Figure S21. Additionally, for the best-performing model, we provide prediction heatmaps showing confidence values for non-decreases only (Suppl. Figure S22) and for both classes combined (Suppl. Figure S23).

**Figure 4:**
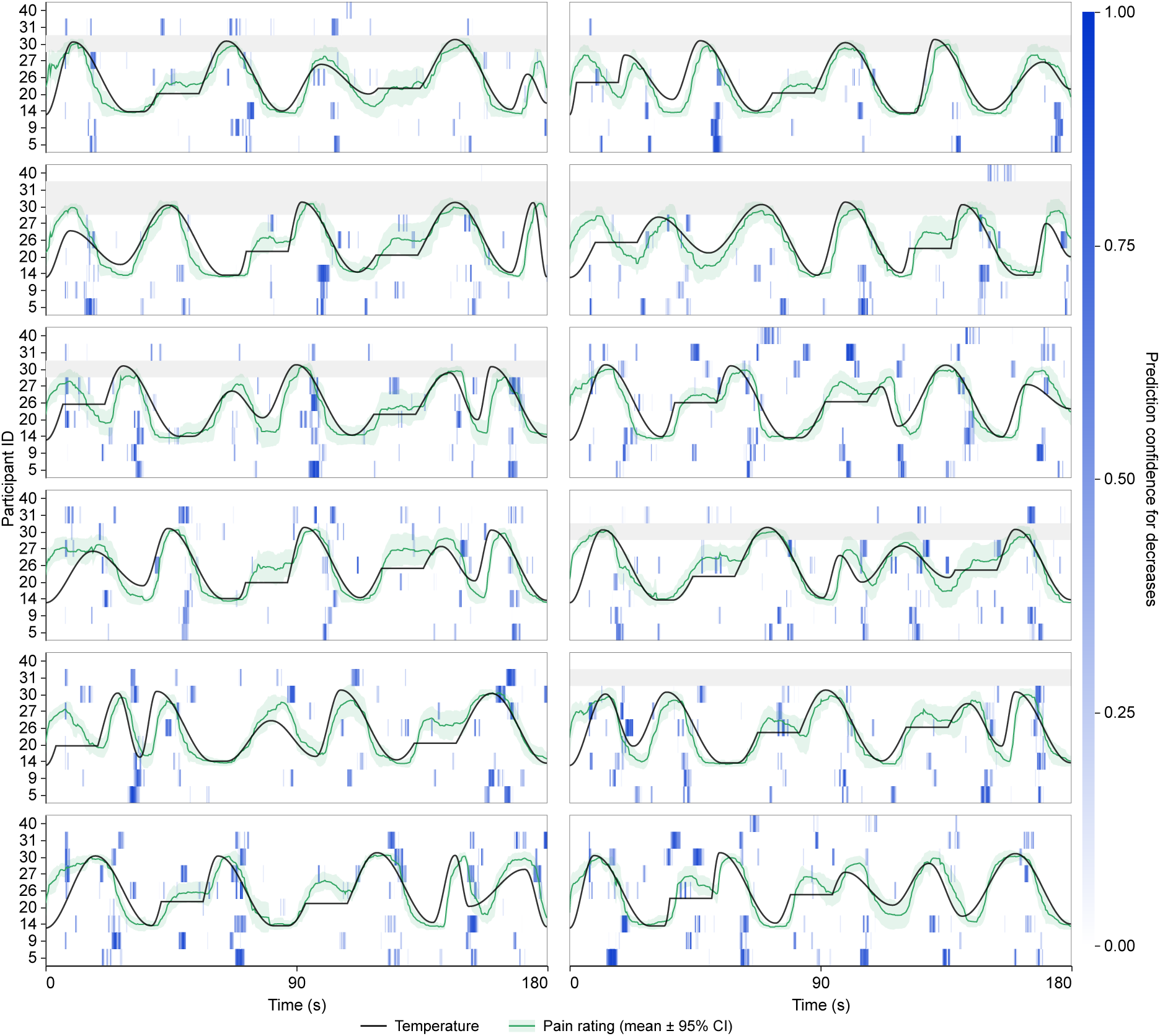
Continuous prediction heatmap for the decrease class using electrodermal activity (EDA), heart rate (HR), and pupil diameter. We applied the trained model using EDA, HR, and pupil diameter to all 12 continuous stimulus curves from the nine held-out participants. Each panel represents one stimulus curve; each heatmap row represents one participant. Blue intensity indicates the model’s predicted probability for the decrease class. Predictions at or above the threshold of 0.90 were classified as decreases. The black line shows the individually calibrated temperature curve; the green line and shaded band show the mean pain rating and 95% confidence interval across the test participants. Grey rows indicate missing trials. Pain ratings were recorded on a visual analogue scale from 0 to 70 and were not used during model training, which relied on temperature intervals as ground truth labels (decreases vs. increases and plateaus) and physiological signals as input. In Suppl. Figure S22 and Figure S23, we additionally provide prediction heatmaps for the detection of non-decreases and for both classes, respectively.

## Discussion

This study tested whether pain decreases can be objectively detected in healthy participants using readily obtainable physiological signals. Results demonstrate that pain decreases can be detected in real-time with substantial accuracy using a combination of electrodermal activity, pupillary responses, and heart rate, with electrodermal activity being the most informative predictor. Despite considerable interindividual variability indicating the need for personalized approaches, these findings provide proof of concept for noninvasive and objective pain monitoring systems.

### Main Findings

#### Detection of Pain Dynamics

This study advances previous research by examining pain dynamics in tonic pain with random fluctuations. Our three-minute noxious heat stimulations fluctuated unpredictably between temperatures calibrated to each participant’s pain threshold and VAS 70. This paradigm more closely approximates chronic pain conditions where individuals experience spontaneous pain fluctuations [7,31,65,66]. Compared with phasic paradigms for pain intensity estimation [26,27,32,35,38,44,59,60,79,81,84] and tonic protocols using constant temperatures [20,68,70,80,82], it allowed us to examine pain decreases during sustained, unpredictable stimulation. By targeting pain changes (first derivative of pain intensity) rather than absolute intensity levels, we may have addressed a more tractable classification problem than absolute intensity estimation. The classifier’s ability to also detect pain decreases that were shorter and based on smaller temperature differences than those used for training further demonstrates generalizability.

#### Multimodal Performance

The combination of electrodermal activity, pupillometry, and heart rate performed best. Electrodermal activity was the most informative single predictor, consistent with its established sensitivity to noxious stimulation through direct sympathetic activation [32,77]. Pupillary responses ranked second, reflecting sympathetic activation through dilator pupillae muscle contraction [22]. Heart rate variability captured sympathico-vagal interactions [63], with noxious stimulation triggering blood pressure increases and subsequent heart rate acceleration [22].

In the model comparison, the PatchTST architecture [69] strikingly outperformed other architectures by over 5 percentage points in accuracy for feature sets containing EDA data (see Suppl. Table S7). The patch-based tokenization of temporal segments aligned optimally with raw EDA data without requiring feature engineering. Our causal preprocessing, which restricted each sample to past and present information, differed from prior work employing non-causal steps for pain intensity estimation [32,35,60,79]. Nevertheless, our minimally processed pipeline achieved strong performance suitable for real-time deployment, validated by continuous-stream analysis showing stable detection during extended periods of fluctuating pain.

These results support the feasibility of predicting pain decreases under causal preprocessing, without being able to attribute the discriminative signal to a specific physiological mechanism. Determining the relative contributions of autonomic dynamics, neural activity, and pain-related behavior will require alternative analyses designed for mechanistic inference that could then also incorporate non-causal preprocessing steps.

#### Individual Variability

Substantial interindividual differences across physiological measures remain critical for clinical translation [22]. Variability differed by modality: EDA and pupillary responses were relatively stable, whereas facial expressions, heart rate, and EEG varied widely with some participants performing even below chance levels, warranting separate discussion.

Facial-expression variability matches Kunz et al. [53], who showed that pain-related action units vary across individuals rather than being uniformly expressed. Additionally, some participants’ pain responses correlated positively with temperature changes, while others showed negative correlations, further complicating universal model development. Facial-expression detection from video may also depend on the specific algorithm [76]. Heart rate showed modest group performance with one notable exception achieving EDA-level accuracy, potentially influenced by breathing patterns under experimental pain [30].

EEG showed the greatest variability, with top and bottom participants similarly distant from chance. This aligns with reports of inconsistent EEG-based pain biomarkers across individuals [12,90,91]. Although enhanced delta and gamma power represent robust neurophysiological findings in pain perception [37,89,91], patterns most relevant for pain decreases remain unclear. Diverse response patterns limited group-level EEG performance and cross-participant generalization. In addition, the minimally preprocessed EEG data for causal real-time settings remain susceptible to non-neural contamination (e.g., EMG-related activity), reducing interpretability [34]. Universal approaches may be limited, making personalized models more appropriate for EEG-based pain detection as shown by Aurucci et al. [4] for the real-time detection of neuropathic pain in patients.

### Limitations & Clinical Translation

#### Generalizability

Generalizability is constrained by several factors. Participants were healthy volunteers with experimentally induced pain; chronic pain patients exhibit distinct EEG signatures and differ in the psychological, social, and pathophysiological dimensions of their pain experience [36,67,87]. The test set comprised nine participants, limiting the precision of performance estimates, though seed stability analysis across multiple splits supported robustness. Financial incentives for accurate pain ratings could potentially have influenced attentional and physiological responses beyond pain perception itself. Nevertheless, establishing proof of concept in a controlled sample provides necessary groundwork for future clinical investigations where physiological heterogeneity may be even greater.

#### Temporal Constraints

Our motivation for this study was to examine whether a device can reliably detect decreases in pain for enhancing treatments such as TENS via precisely timed interventions. Once a decrease is identified, the device would offer the participant a treatment option, an option that would be perceived as effective because the pain is already declining. This could re-establish a sense of control despite the reversed causal order (the decrease is detected first, and the treatment follows). Through the illusion of control, this sequence may nevertheless be experienced as a genuine instance of self-initiated pain relief [16].

Timing is crucial: patients must be offered a treatment option as early as possible during a spontaneous pain decrease, allowing them to attribute the relief to their own actions rather than to uncontrollable fluctuations. This requires minimal detection delays to preserve temporal coupling between a potential treatment and pain relief. Given EDA’s role as the primary predictor, sweat-gland response times impose a physiological delay for the detection of pain changes [13]. In our data, phasic EDA lagged temperature by approximately 2.5 seconds in cross-correlation analysis, and pain decreases were detected at a median of 5.75 seconds.

In an unreported analysis, shorter sample windows (4-6 s) reduced accuracy without improving detection speed.

#### Classification Accuracy

Our models achieved reasonable accuracy with false-positive rates varying across participants and measures. False positives occurred mainly during temperature plateaus. However, these apparent misclassifications as decreases often coincided with subjective pain decreases (see Figure 4), suggesting detection of habituation-related reductions rather than simple tracking of temperature labels. Classifier performance might improve if labels were derived from pain ratings rather than temperature profiles.

Notably, these classification errors need not undermine therapeutic efficacy. False positive detections would result in participants administering a treatment during moments when pain does not decrease. If, however, other actual pain decreases are correctly detected, this could lead to a partial reinforcement schedule, which could produce more extinction-resistant learning than continuous reinforcement [28], potentially enhancing expectation effects [86]. This would also allow for less strict classification thresholds with the additional benefit of lower latencies by prioritizing sensitivity over specificity. Moreover, cue-relief associations could first be established through prior conditioning with experimentally induced pain, where covert reductions in nociceptive input reliably follow the treatment cue, as such pairings have been shown to confer genuine expectation-based analgesic properties [46,71].

Although multimodal ensembles theoretically outperform single modalities through noise reduction, results revealed nuanced patterns. The addition of EDA and pupil diameter signals consistently improved performance, whereas facial expressions, heart rate, and EEG provided no benefits. Full ensemble performance was inferior to optimal subsets, suggesting trade-offs between information gain and noise accumulation, and supporting personalized, fine-tuned approaches consistent with precision medicine principles [42].

### Future Implementation

Results indicate that while the best-performing pain detection model combines EDA, HR, and pupil signals, conceivably enhanced by personalized, fine-tuned EEG and facial expression data, the most pragmatic approach uses only EDA and HR measurements. Despite high univariate pupillometry performance, real-world implementation encounters constraints from ambient light variations and screen usage. Although EEG could potentially improve latency and accuracy, its application has several drawbacks, namely a high susceptibility to noise and movement artifacts, the need for fine-tuned models, and the impractical equipment required for measurement. Facial expression analysis faces similar challenges with specialized cameras or electrodes. EDA and heart rate, on the other hand, can easily be implemented using commercially available, wearable devices [5,19,45]. In the domain of pain versus no-pain detection, previous work has demonstrated feasibility for the real-time detection of experimental pain in healthy participants using EDA only [6,47], further supporting this pragmatic approach.

In a practical implementation, a wrist-worn device would use heart-rate thresholds and accelerometry to restrict detection to sedentary, low-motion periods. Once a high-confidence pain decrease is detected based on EDA and HR, the wrist-device or a smartphone application could prompt the patient to initiate a brief intervention such as TENS via a simple button press. The therapeutic rationale is the timing: because pain is already decreasing, the self-initiated TENS treatment should be experienced as the cause of the (already ongoing) relief and thus strengthen perceived control and self-efficacy through repeated association.

This study demonstrated that the objective, near real-time detection of pain decreases is feasible using easily obtainable physiological signals in healthy participants. Clinical validation remains essential for realizing the therapeutic potential of these findings in preventing pain chronification through precisely timed interventions.

## Acknowledgements

The authors thank all participants for participating in the study. This work was supported by the European Research Council (ERC) under the grant ERC-AdG-883892-PainPersist. The authors have no conflicts of interest to declare. AI tools (Claude, ChatGPT) assisted with manuscript editing and code generation. The authors reviewed and take full responsibility for all content. All data will be made available in a forthcoming data publication. In the meantime, data will be available upon reasonable request to the corresponding author. All code is publicly available at: https://github.com/visserle/pain-measurement.

## Supplementary Materials

### Supplementary Material A: Experimental Design and Procedures

**Figure S1:**
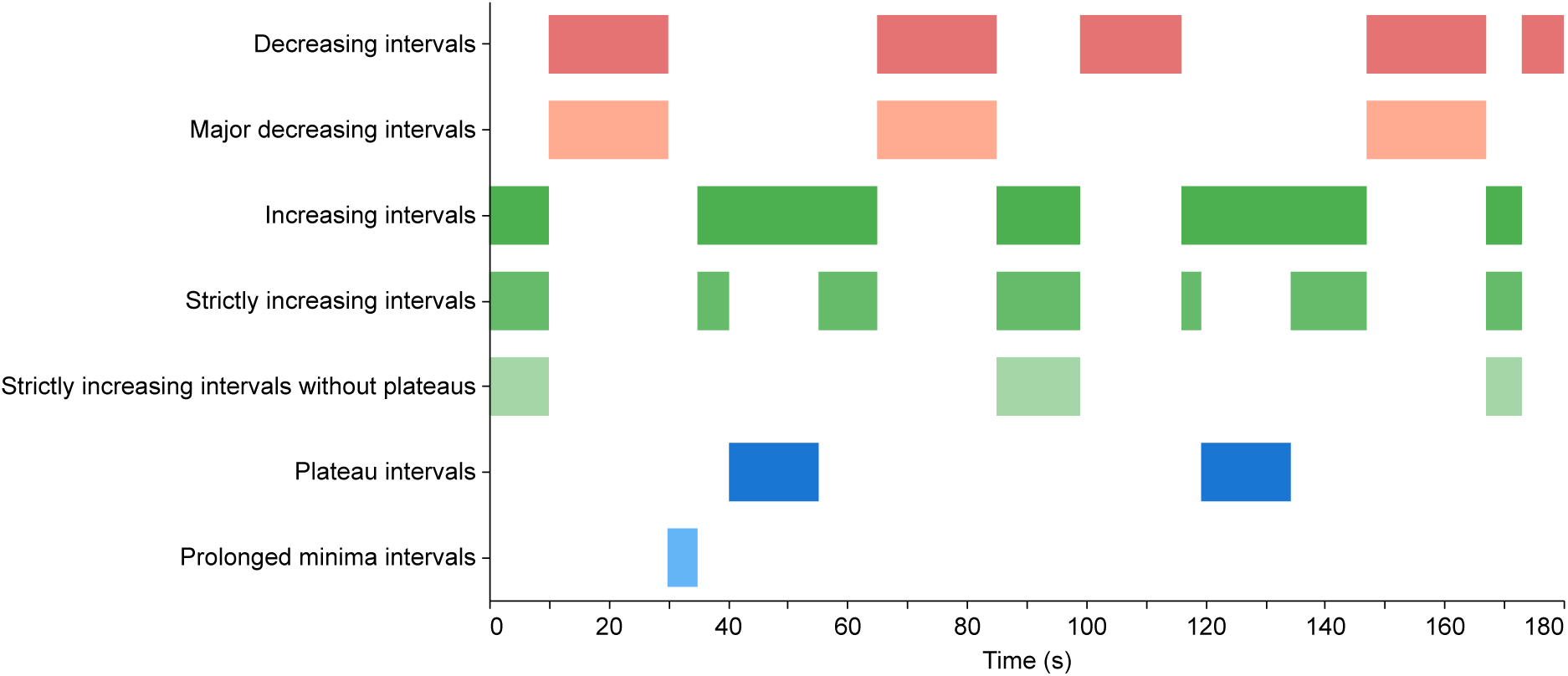
Stimulus interval types used to construct classification samples. Colored bars mark the interval labels derived automatically from the example stimulus curve generated with random seed 133. Samples for the classification task were based on strictly increasing intervals without plateaus, strictly increasing intervals after plateaus, and plateau intervals for non-decreases, and major decreasing intervals for decreases.

**Figure S2:**
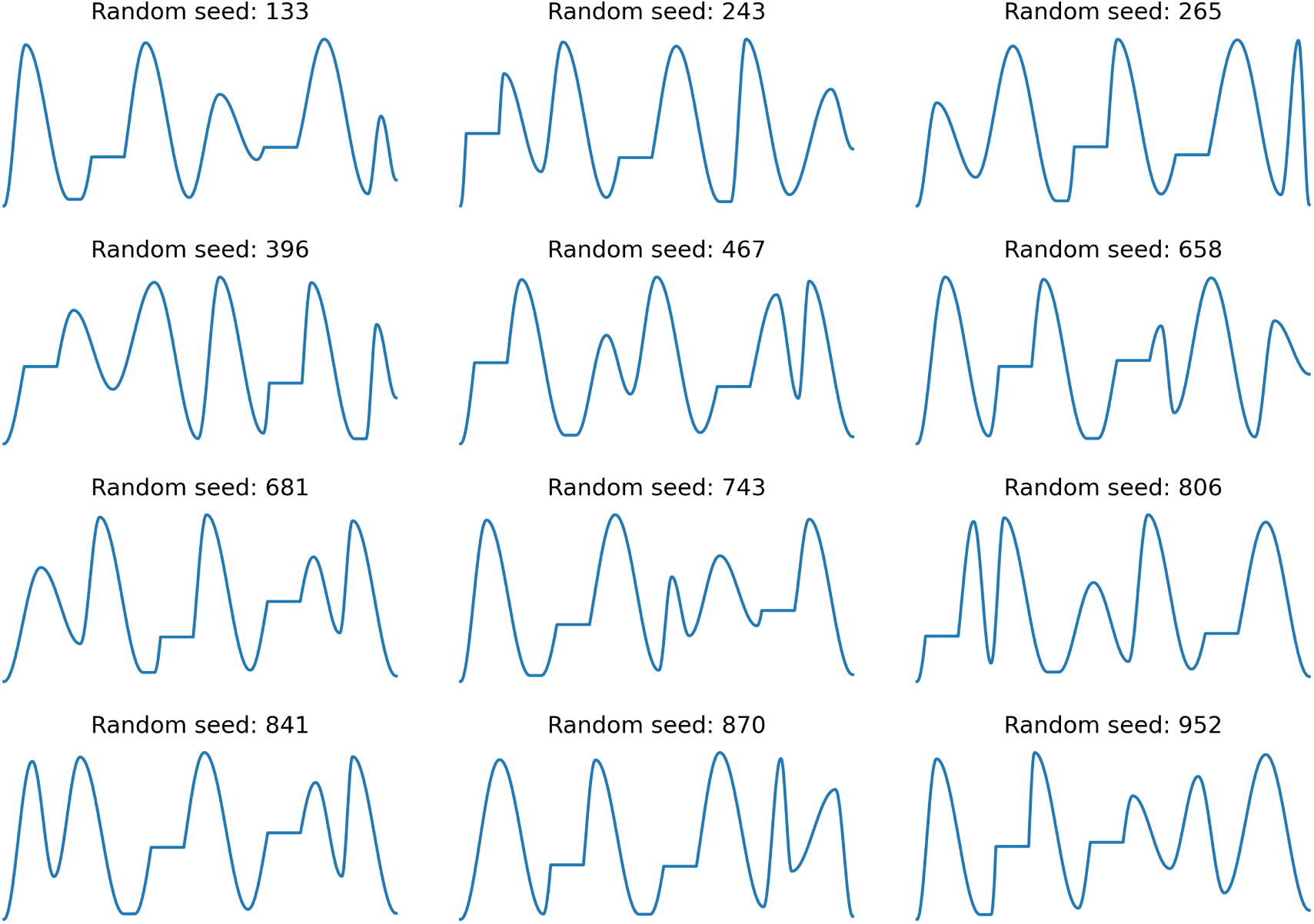
Temperature curves used in the pain measurement experiment. The random seeds determined the output of the stimulus generator.

**Figure S3:**
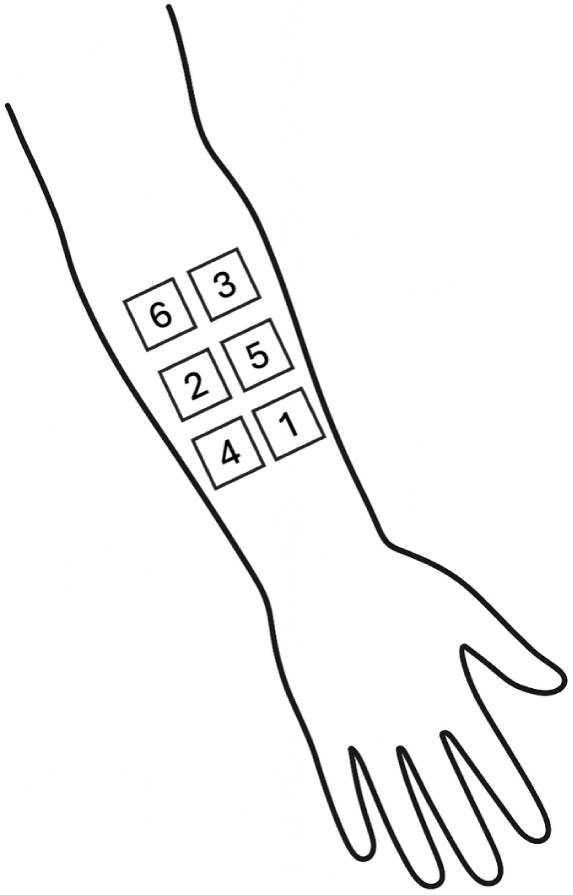
Distribution of skin patches. Six patches on the left inner forearm were arranged in two parallel columns (three per column). To minimize habituation and sensitization effects, we alternated between patches to maximize the distance between two consecutive stimulations. Throughout the experiment, each patch received two stimulations. Stimulation order was counterbalanced across participants: participants with odd IDs received stimulation in ascending order (1-6, repeated twice) with calibration on patch 5; even IDs received descending order (6-1, repeated twice) with calibration on patch 2.

### Supplementary Material B: Participant Characteristics

**Table S1:** Demographic characteristics of all participants.

| 1059 | ID | Age | Gender | BMI | Education | Employment | Physical activity | Meditation |
| --- | --- | --- | --- | --- | --- | --- | --- | --- |
| 1060 | 1 | 22 | f | 22 | University degree | Student | Several times a week | No |
| 1061 | 2 | 23 | f | 27 | Abitur (university entrance qualification) | Student | Daily | No |
| 1062 | 3 | 23 | f | 23 | University degree | Employed part-time | Several times a week | No |
| 1063 | 4 | 35 | f | 20 | University degree | Unemployed | Several times a week | No |
| 1064 | 5 | 25 | f | 27 | Trade/technical/vocational training | Student | Several times a week | No |
| 1065 | 6 | 20 | f | 23 | Abitur (university entrance qualification) | Student | Several times a week | No |
| 1066 | 7 | 27 | f | 19 | Abitur (university entrance qualification) | Student | Several times a week | No |
| 1067 | 8 | 23 | f | 20 | Abitur (university entrance qualification) | Student | Several times a week | No |
| 1068 | 9 | 23 | f | 27 | Abitur (university entrance qualification) | Student | Once a week | No |
| 1069 | 10 | 22 | f | 21 | Abitur (university entrance qualification) | Student | Several times a week | Yes |
| 1070 | 11 | 18 | f | 20 | Abitur (university entrance qualification) | Student | Several times a week | No |
| 1071 | 12 | 22 | f | 21 | Abitur (university entrance qualification) | Student | Once a week | No |
| 1072 | 13 | 39 | f | 26 | Secondary school leaving certificate (Mittlere Reife) | Employed full-time | Once a week | No |
| 1073 | 14 | 30 | f | 22 | Abitur (university entrance qualification) | Student | A few times a month | No |
| 1074 | 15 | 23 | f | 28 | Abitur (university entrance qualification) | Student | Once a week | No |
| 1075 | 16 | 29 | f | 29 | Abitur (university entrance qualification) | Unemployed | A few times a month | No |
| 1076 | 17 | 26 | f | 21 | University degree | Student | Several times a week | No |
| 1077 | 18 | 24 | f | 26 | Abitur (university entrance qualification) | Student | Several times a week | No |
| 1078 | 19 | 27 | f | 20 | Abitur (university entrance qualification) | Student | Several times a week | No |
| 1079 | 20 | 27 | f | 22 | Abitur (university entrance qualification) | Student | Daily | No |
| 1080 | 21 | 22 | f | 20 | University degree | Student | Several times a week | No |
| 1081 | **ID** | **Age** | **Gender** | **BMI** | **Education** | **Employment** | **Physical activity** | **Meditation** |
| 1082 | 22 | 27 | f | 24 | University degree | Employed part-time | Once a week | No |
| 1083 | 23 | 27 | f | 23 | Trade/technical/vocational training | Student | A few times a month | No |
| 1084 | 24 | 22 | f | 23 | Abitur (university entrance qualification) | Student | Several times a week | No |
| 1085 | 25 | 25 | f | 21 | Abitur (university entrance qualification) | Student | Several times a week | No |
| 1086 | 26 | 27 | f | 21 | University degree | Employed full-time | Several times a week | Yes |
| 1087 | 27 | 34 | f | 22 | University degree | Student | Several times a week | No |
| 1088 | 28 | 22 | f | 26 | Abitur (university entrance qualification) | Student | Several times a week | No |
| 1089 | 29 | 22 | f | 26 | Abitur (university entrance qualification) | Student | Several times a week | No |
| 1090 | 30 | 34 | f | 24 | Abitur (university entrance qualification) | Student | Several times a week | No |
| 1091 | 31 | 28 | f | 26 | University degree | Unemployed | Several times a week | No |
| 1092 | 32 | 26 | f | 20 | University degree | Student | Several times a week | No |
| 1093 | 33 | 22 | f | 21 | Abitur (university entrance qualification) | Student | Once a week | No |
| 1094 | 34 | 25 | f | 25 | University degree | Student | Several times a week | No |
| 1095 | 35 | 20 | f | 20 | Abitur (university entrance qualification) | Student | Several times a week | No |
| 1096 | 36 | 39 | f | 27 | Trade/technical/vocational training | Student | A few times a month | No |
| 1097 | 37 | 35 | f | 23 | University degree | Student | A few times a month | No |
| 1098 | 38 | 24 | f | 25 | Abitur (university entrance qualification) | Student | Once a week | No |
| 1099 | 39 | 25 | f | 20 | University degree | Student | Several times a week | No |
| 1100 | 40 | 35 | f | 22 | University degree | Unemployed | A few times a month | No |
| 1101 | 41 | 28 | f | 30 | Trade/technical/vocational training | Student | Several times a week | No |
| 1102 | 42 | 22 | f | 28 | Abitur (university entrance qualification) | Student | Several times a week | No |

*Note:* ID = anonymized participant identifier; BMI = body mass index (kg/m<sup>2</sup>); Physical activity = exercise frequency; Meditation = self-reported practice (yes/no).

**Table S2:** BDI-II and STAI-T-10 scores and PCS total and subscale scores.

| ID | BDI-II | STAI-T-10 | PCS<br>Total | PCS<br>Rumination | PCS<br>Magnification | PCS<br>Helplessness |
| --- | --- | --- | --- | --- | --- | --- |
| 1 | 1 | 18.57 | 14 | 7 | 5 | 2 |
| 2 | 8 | 34.29 | 26 | 10 | 6 | 10 |
| 3 | 3 | 38.57 | 17 | 7 | 2 | 8 |
| 4 | 0 | 68.57 | 26 | 13 | 3 | 10 |
| 5 | 4 | 17.14 | 19 | 9 | 4 | 6 |
| 6 | 1 | 45.71 | 17 | 6 | 4 | 7 |
| 7 | 0 | 48.57 | 20 | 8 | 3 | 9 |
| 8 | 3 | 30 | 22 | 7 | 6 | 9 |
| 9 | 1 | 32.86 | 20 | 7 | 5 | 8 |
| 10 | 5 | 35.71 | 29 | 8 | 9 | 12 |
| 11 | 3 | 38.57 | 9 | 6 | 0 | 3 |
| 12 | 2 | 62.86 | 12 | 1 | 3 | 8 |
| 13 | 3 | 30 | 13 | 4 | 2 | 7 |
| 14 | 0 | 2.86 | 16 | 8 | 1 | 7 |
| 15 | 6 | 68.57 | 18 | 9 | 4 | 5 |
| 16 | 4 | 38.57 | 32 | 13 | 6 | 13 |
| 17 | 3 | 42.86 | 16 | 4 | 6 | 6 |
| 18 | 5 | 22.86 | 28 | 13 | 4 | 11 |
| 19 | 0 | 20 | 30 | 11 | 5 | 14 |
| 20 | 0 | 15.71 | 7 | 1 | 0 | 6 |
| 21 | 0 | 5.71 | 1 | 0 | 1 | 0 |
| 22 | 4 | 45.71 | 17 | 7 | 6 | 4 |
| 23 | 4 | 41.43 | 23 | 8 | 8 | 7 |
| 24 | 1 | 21.43 | 0 | 0 | 0 | 0 |
| 25 | 0 | 55.71 | 33 | 13 | 8 | 12 |
| 26 | 0 | 14.29 | 14 | 8 | 3 | 3 |
| 27 | 0 | 41.43 | 28 | 10 | 5 | 13 |
| 28 | 9 | 48.57 | 20 | 13 | 0 | 7 |
| 29 | 1 | 27.14 | 21 | 11 | 4 | 6 |
| 30 | 0 | 7.14 | 2 | 1 | 0 | 1 |
| 31 | 0 | 12.86 | 0 | 0 | 0 | 0 |
| 32 | 12 | 57.14 | 16 | 9 | 6 | 1 |
| 33 | 10 | 35.71 | 16 | 6 | 4 | 6 |
| 34 | 7 | 38.57 | 24 | 13 | 4 | 7 |
| 35 | 8 | 51.43 | 25 | 9 | 6 | 10 |
| 36 | 3 | 25.71 | 19 | 8 | 4 | 7 |
| 37 | 0 | 31.43 | 0 | 0 | 0 | 0 |
| 38 | 7 | 47.14 | 22 | 9 | 6 | 7 |
| 39 | 4 | 25.71 | 21 | 9 | 4 | 8 |
| 40 | 1 | 48.57 | 9 | 4 | 2 | 3 |
| 41 | 11 | 35.71 | 20 | 8 | 3 | 9 |
| 42 | 10 | 48.57 | 15 | 2 | 5 | 8 |

*Note:* BDI-II = Beck Depression Inventory II; STAI-T-10 = 10-item State-Trait Anxiety Inventory, Trait version; PCS = Pain Catastrophizing Scale. Total denotes the overall PCS score; Rumination, Magnification, and Helplessness are PCS subscale scores.

**Table S3:** Mindful Attention Awareness Scale (MAAS) and Pain Vigilance and Awareness Questionnaire (PVAQ) scores.

| ID | MAAS | PVAQ<br>Total | PVAQ<br>Attention to pain | PVAQ<br>Attention to changes |
| --- | --- | --- | --- | --- |
| 1 | 4.53 | 37 | 19 | 18 |
| 2 | 3.67 | 37 | 20 | 17 |
| 3 | 4.13 | 23 | 12 | 11 |
| 4 | 3.33 | 44 | 21 | 23 |
| 5 | 3.2 | 32 | 16 | 16 |
| 6 | 4 | 37 | 23 | 14 |
| 7 | 4.6 | 38 | 18 | 20 |
| 8 | 3.47 | 36 | 18 | 18 |
| 9 | 4.27 | 35 | 20 | 15 |
| 10 | 3 | 53 | 31 | 22 |
| 11 | 4.2 | 30 | 10 | 20 |
| 12 | 3.2 | 33 | 17 | 16 |
| 13 | 3.47 | 46 | 24 | 22 |
| 14 | 4.87 | 32 | 16 | 16 |
| 15 | 3.47 | 39 | 20 | 19 |
| 16 | 3.73 | 45 | 22 | 23 |
| 17 | 4.2 | 28 | 13 | 15 |
| 18 | 4.87 | 60 | 35 | 25 |
| 19 | 4.67 | 41 | 21 | 20 |

| ID | MAAS | PVAQ | PVAQ | PVAQ |
| --- | --- | --- | --- | --- |
|  |  | Total | Attention to pain | Attention to changes |
| 20 | 5.53 | 31 | 15 | 16 |
| 21 | 5.4 | 54 | 27 | 27 |
| 22 | 3.53 | 51 | 31 | 20 |
| 23 | 4.6 | 27 | 10 | 17 |
| 24 | 4.47 | 37 | 21 | 16 |
| 25 | 4 | 42 | 25 | 17 |
| 26 | 3.93 | 30 | 16 | 14 |
| 27 | 4.2 | 50 | 28 | 22 |
| 28 | 2.93 | 33 | 12 | 21 |
| 29 | 3.93 | 34 | 18 | 16 |
| 30 | 4.8 | 35 | 21 | 14 |
| 31 | 3.27 | 28 | 16 | 12 |
| 32 | 3.93 | 49 | 26 | 23 |
| 33 | 3.6 | 29 | 14 | 15 |
| 34 | 4.33 | 24 | 9 | 15 |
| 35 | 3.47 | 40 | 23 | 17 |
| 36 | 5.33 | 46 | 25 | 21 |
| 37 | 4 | 51 | 27 | 24 |
| 38 | 3 | 46 | 26 | 20 |
| 39 | 3.6 | 37 | 18 | 19 |
| 40 | 5.6 | 45 | 20 | 25 |
| 41 | 3.47 | 31 | 12 | 19 |
| 42 | 1.47 | 29 | 15 | 14 |

*Note:* MAAS = Mindful Attention Awareness Scale; PVAQ = Pain Vigilance and Awareness Questionnaire. Total denotes the overall PVAQ score; Attention to pain and Attention to changes are PVAQ subscale scores.

**Table S4:**
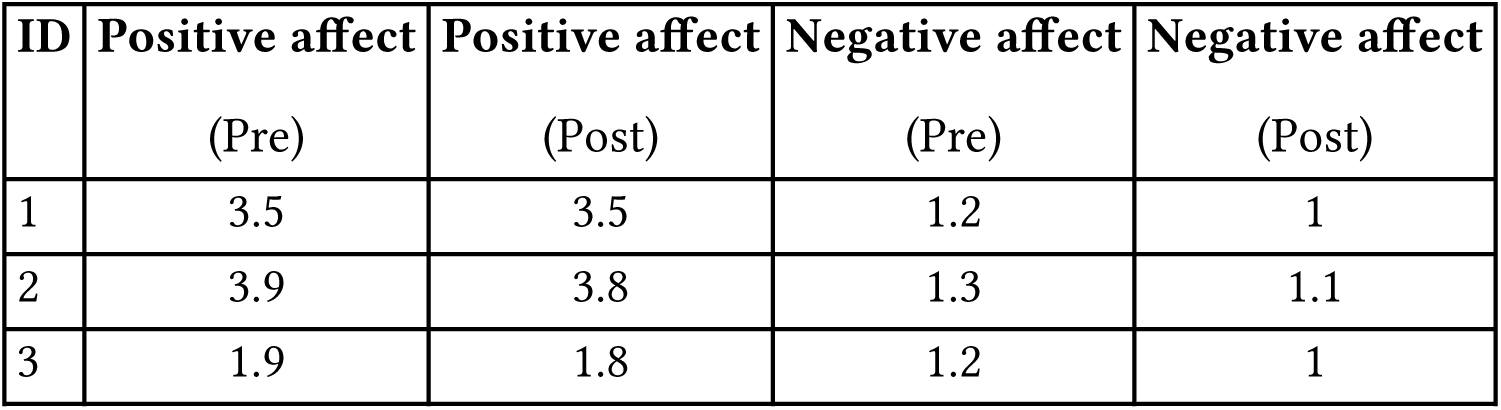
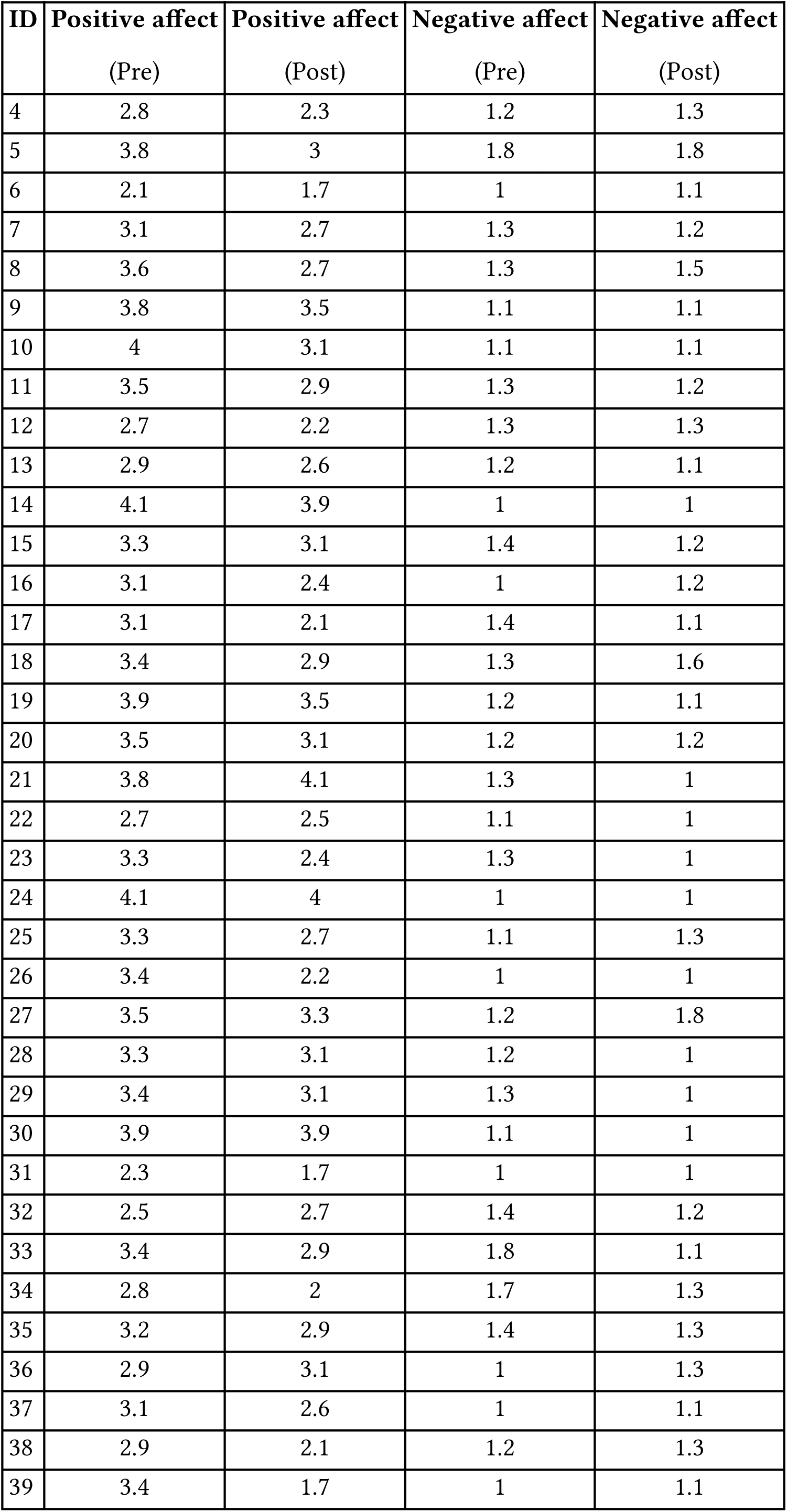
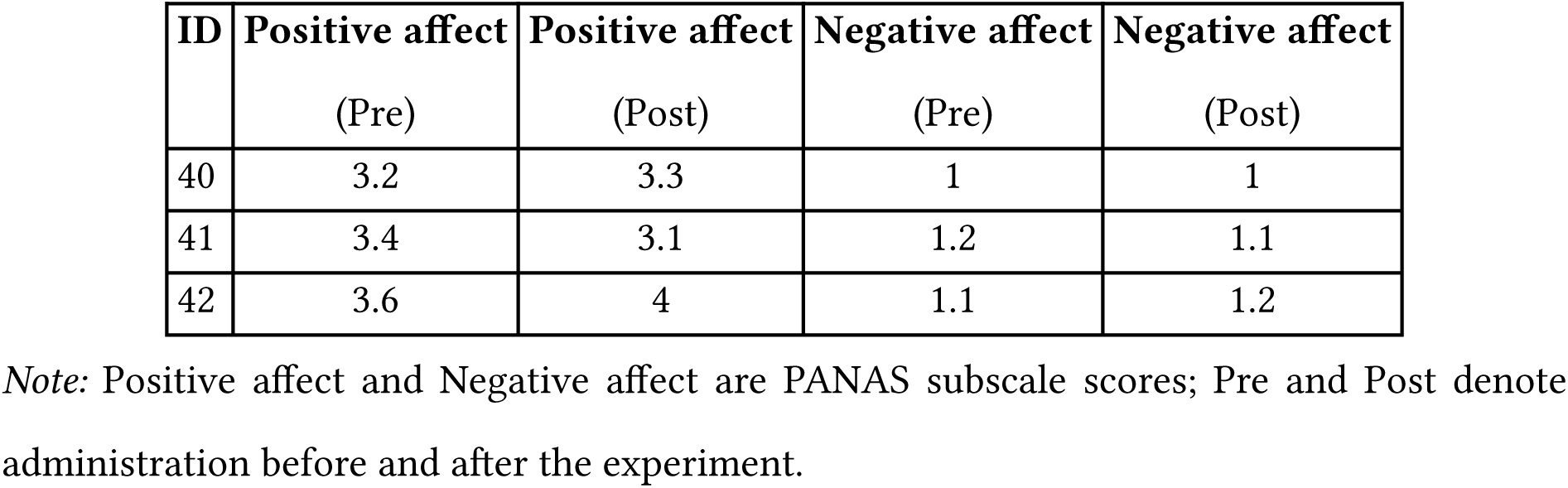
Positive and Negative Affect Schedule (PANAS) scores before and after the experiment.

| ID | Positive affect | Positive affect | Negative affect | Negative affect |
| --- | --- | --- | --- | --- |
|  | (Pre) | (Post) | (Pre) | (Post) |
| 1 | 3.5 | 3.5 | 1.2 | 1 |
| 2 | 3.9 | 3.8 | 1.3 | 1.1 |
| 3 | 1.9 | 1.8 | 1.2 | 1 |

|  |  |  |  |  |  |
| --- | --- | --- | --- | --- | --- |
| 1213 | **ID** | **Positive affect** | **Positive affect** | **Negative affect** | **Negative affect** |
| 1214 |  | (Pre) | (Post) | (Pre) | (Post) |
| 1215 | 4 | 2.8 | 2.3 | 1.2 | 1.3 |
| 1216 | 5 | 3.8 | 3 | 1.8 | 1.8 |
| 1217 | 6 | 2.1 | 1.7 | 1 | 1.1 |
| 1218 | 7 | 3.1 | 2.7 | 1.3 | 1.2 |
| 1219 | 8 | 3.6 | 2.7 | 1.3 | 1.5 |
| 1220 | 9 | 3.8 | 3.5 | 1.1 | 1.1 |
| 1221 | 10 | 4 | 3.1 | 1.1 | 1.1 |
| 1222 | 11 | 3.5 | 2.9 | 1.3 | 1.2 |
| 1223 | 12 | 2.7 | 2.2 | 1.3 | 1.3 |
| 1224 | 13 | 2.9 | 2.6 | 1.2 | 1.1 |
| 1225 | 14 | 4.1 | 3.9 | 1 | 1 |
| 1226 | 15 | 3.3 | 3.1 | 1.4 | 1.2 |
| 1227 | 16 | 3.1 | 2.4 | 1 | 1.2 |
| 1228 | 17 | 3.1 | 2.1 | 1.4 | 1.1 |
| 1229 | 18 | 3.4 | 2.9 | 1.3 | 1.6 |
| 1230 | 19 | 3.9 | 3.5 | 1.2 | 1.1 |
| 1231 | 20 | 3.5 | 3.1 | 1.2 | 1.2 |
| 1232 | 21 | 3.8 | 4.1 | 1.3 | 1 |
| 1233 | 22 | 2.7 | 2.5 | 1.1 | 1 |
| 1234 | 23 | 3.3 | 2.4 | 1.3 | 1 |
| 1235 | 24 | 4.1 | 4 | 1 | 1 |
| 1236 | 25 | 3.3 | 2.7 | 1.1 | 1.3 |
| 1237 | 26 | 3.4 | 2.2 | 1 | 1 |
| 1238 | 27 | 3.5 | 3.3 | 1.2 | 1.8 |
| 1239 | 28 | 3.3 | 3.1 | 1.2 | 1 |
| 1240 | 29 | 3.4 | 3.1 | 1.3 | 1 |
| 1241 | 30 | 3.9 | 3.9 | 1.1 | 1 |
| 1242 | 31 | 2.3 | 1.7 | 1 | 1 |
| 1243 | 32 | 2.5 | 2.7 | 1.4 | 1.2 |
| 1244 | 33 | 3.4 | 2.9 | 1.8 | 1.1 |
| 1245 | 34 | 2.8 | 2 | 1.7 | 1.3 |
| 1246 | 35 | 3.2 | 2.9 | 1.4 | 1.3 |
| 1247 | 36 | 2.9 | 3.1 | 1 | 1.3 |
| 1248 | 37 | 3.1 | 2.6 | 1 | 1.1 |
| 1249 | 38 | 2.9 | 2.1 | 1.2 | 1.3 |
| 1250 | 39 | 3.4 | 1.7 | 1 | 1.1 |

| ID | Positive affect<br>(Pre) | Positive affect<br>(Post) | Negative affect<br>(Pre) | Negative affect<br>(Post) |
| --- | --- | --- | --- | --- |
| 40 | 3.2 | 3.3 | 1 | 1 |
| 41 | 3.4 | 3.1 | 1.2 | 1.1 |
| 42 | 3.6 | 4 | 1.1 | 1.2 |

*Note:* Positive affect and Negative affect are PANAS subscale scores; Pre and Post denote administration before and after the experiment.

### Supplementary Material C: Deep Learning Model Architectures

#### Multilayer Perceptron

A multilayer perceptron (MLP) is a feedforward artificial neural network consisting of an input layer, one or more hidden layers, and an output layer. Information flows unidirectionally from the input layer through the hidden layers to the output layer. Each layer computes a weighted sum of its inputs followed by a nonlinear activation function (commonly ReLU or sigmoid). During training, the network optimizes its weights and biases through backpropagation, an algorithm that iteratively adjusts parameters to minimize a loss function by computing gradients via the chain rule. MLPs are universal function approximators [4] capable of learning complex nonlinear relationships between input features and target outputs, making them well-suited for classifications. For multivariate time series, the input is flattened into a single input vector (1D) as MLPs inherently do not understand temporal structure [7].

#### LightTS

LightTS is a lightweight MLP-based architecture designed for efficient multivariate time series processing [14]. It implements two complementary down-sampling strategies before applying the data to MLP layers: interval sampling and continuous sampling. Interval sampling extracts values at regular intervals to capture long-term patterns, while continuous sampling selects contiguous subsequences to preserve local temporal dependencies. These down-sampled representations are then processed through separate MLP structures, and their outputs are aggregated to produce the final output. This design exploits the observation that down-sampled time series often retain the majority of relevant information while drastically reducing computational requirements [14].

#### TimesNet

TimesNet is an architecture based on convolutional neural networks (CNNs) and it leverages the fact that many time series exhibit multiple periodic patterns [13]. Rather than processing temporal data in its native 1D format, TimesNet first identifies dominant periods in the signal and reshapes the 1D sequence into 2D tensors according to these periods. This transformation arranges variations within each cycle as columns and variations across cycles as rows, essentially converting temporal patterns into spatial patterns. The architecture then applies special 2D convolutional kernels (inception blocks) to extract features from these reshaped representations. This approach is more parameter-efficient than modeling complex temporal dependencies directly in 1D. In the review by Wang et al. [12], this architecture achieved the best performance for classification tasks across a variety of multivariate time series compared with other state-of-the-art deep learning architectures.

#### PatchTST

PatchTST’s architecture is based on the transformer architecture and is designed for the efficient analysis of multivariate time series [10]. Instead of treating individual time points as tokens (discrete input units processed by the model), it segments the input sequence into patches that serve as input to the transformer encoder. PatchTST employs a channel-independent strategy where each univariate series is processed separately with shared weights. This approach reduces both overfitting risk and computational requirements. The self-attention mechanism (a method that weighs the relevance of different parts of the input to each other) captures long-range dependencies between patches. This makes PatchTST particularly suitable for multivariate physiological signal processing where multiple channels need to be analyzed efficiently.

#### iTransformer

iTransformer is a transformer-based architecture that inverts the traditional application of attention mechanisms in multivariate time series analysis [9]. Unlike conventional transformers that apply attention across time points, iTransformer treats each variate (each individual series within the multivariate dataset) as a token and applies self-attention across the variate dimension. This inversion allows the model to capture multivariate correlations directly while reducing computational complexity. Conceptually, this resembles a learned, dynamic correlation matrix that adapts to the input data. The feed-forward network is applied to each variate token to learn nonlinear representations. The design is particularly advantageous when inter-variate relationships are more informative than temporal patterns, such as in physiological signal analysis where multiple biosignals interact simultaneously.

#### EEGNet

EEGNet is an efficient convolutional neural network architecture specifically designed for EEG-based brain-computer interfaces [6]. It incorporates domain knowledge from EEG analysis by employing temporal convolution to learn frequency-specific filters, depthwise spatial convolution to capture optimal electrode combinations, and separable convolutions to extract additional features in the data. Despite its compact size, EEGNet demonstrates competitive performance across diverse EEG classification tasks and performs well even when training data is limited [6].

#### Ensemble Models

The ensemble models combined EEGNet for EEG data and PatchTST for peripheral physio-logical and facial expression data. This design grouped related signals and addressed the different sampling rates inherent in our multimodal dataset: EEG signals at 250 Hz and physiological signals (EDA, heart rate, pupil diameter, facial expressions) at 10 Hz. For feature sets combining EEG with other physiological data, non-EEG signals were upsampled to 250 Hz using linear interpolation to match the EEG sampling rate and to enable simple sample generation in PyTorch. The architecture split the input by channel, routing EEG channels through EEGNet and other channels through PatchTST. The upsampled input for PatchTST was downsampled back to 10 Hz (taking every 25th data point). Each sub-model had its final classification layers removed and instead served as feature extractors. The extracted features were concatenated and passed through a fusion network consisting of MLP layers. This feature-level fusion approach [2] allowed each sub-model to learn modality-specific representations optimally before combining them for classification, a strategy shown effective for multimodal physiological signal processing [3].

For non-ensemble models, all upsampled signals were fed directly alongside EEG as a unified input with a sampling rate of 250 Hz.

### Supplementary Material D: Hyperparameter Optimization, Model

#### Training and Evaluation

##### Hyperparameter Optimization

We conducted hyperparameter optimization using the Optuna library (version 4.3.0) [1], which implements Bayesian-inspired optimization to efficiently search the hyperparameter space. The hyperparameter search spaces (see Suppl. Table S8) were informed by default values from Wang et al. [12] and Lawhern et al. [6]. We further included learning rate as a tuneable hyperparameter (range: 0.00001 to 0.01, log-scaled), given its critical influence on convergence and performance [8]. For 30 trials, each model’s hyperparameter space was optimized by maximizing the accuracy of the validation set. A single trial consisted of 15 epochs with a batch size of 64. We used Optuna-based median pruning (5 startup trials, 5 warmup steps) to terminate unpromising trials early. Additionally, we used a fixed learning rate decay schedule (cosine annealing without restarts and with a maximum number of iterations equaling the number of epochs) to eliminate dependence on validation monitoring during the final model training.

#### Model Training

We employed cross-entropy as the loss function for our binary classification task. Cross-entropy is defined as

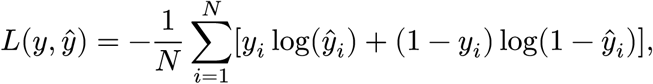

where *y* represents the vector of true labels, 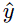 is the vector of predicted probabilities, and *N* is the number of samples in each batch. For optimization, we utilized the Adam optimizer [5]. To prevent exploding gradients and ensure stable training, we applied gradient clipping with a maximum norm of 4.0.

Model training was performed using PyTorch (version 2.6.0) [11] on an NVIDIA GeForce RTX 3080 Ti GPU. We set a fixed random seed (42) across all experiments for Python, NumPy, PyTorch and Optuna to ensure reproducibility of results. The complete hyperparameter optimization cycle took approximately 3 hours for all model-feature combinations.

#### Model Evaluation

We utilized accuracy as our primary evaluation metric during model selection. Following this, we evaluated the best-performing model for each feature set using additional classification metrics: area under the receiver operating characteristic curve (AUROC), precision, recall, *F*_1_ – scores and Matthews correlation coefficient (MCC). We defined pain decreases as positive and non-decreases as negative cases. Let TP, TN, FP, and FN denote true positives, true negatives, false positives, and false negatives, respectively. Accuracy quantified the overall proportion of correct predictions:

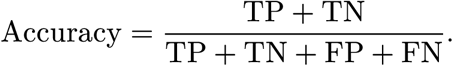

AUROC quantified the model’s ability to discriminate between pain decreases and non-decreases across all possible classification thresholds. This metric ranges from 0 to 1, where values near 0.5 indicate chance-level performance, and scores approaching 1 demonstrate strong discriminative capability. Precision (also called positive predictive value (PPV)) evaluated the proportion of true positives among predicted positives, indicating the reliability of positive predictions. Recall (also known as sensitivity) measured the model’s ability to correctly identify positive classes (pain decreases), representing the proportion of actual positives correctly identified by the model. The F₁-score provided the harmonic mean of precision and recall, offering a balance between these two metrics particularly useful when class distributions are imbalanced:

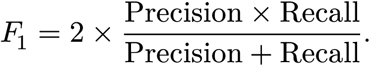

MCC quantified the association between observed and predicted classifications, producing a value between −1 and +1, where +1 represents perfect prediction, 0 represents random prediction, and −1 represents inverse prediction:

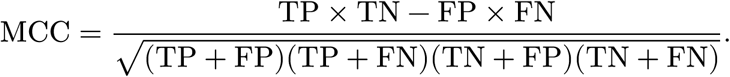

### Supplementary Material E: Exploratory Data Analysis

#### Psychophysiological Signals

**Figure S4:**
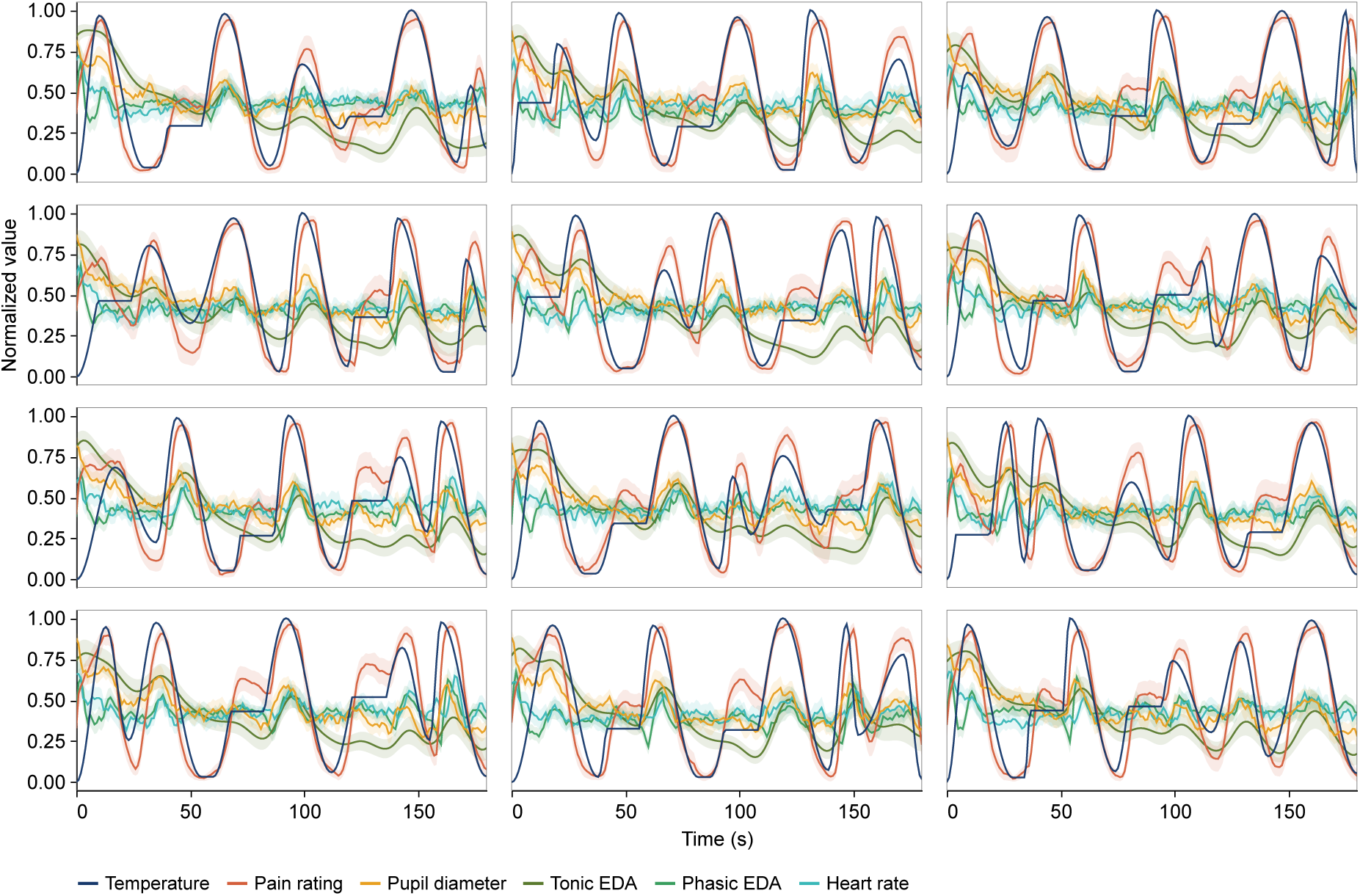
Grand-averaged means of physiological signals.

**Figure S5:**
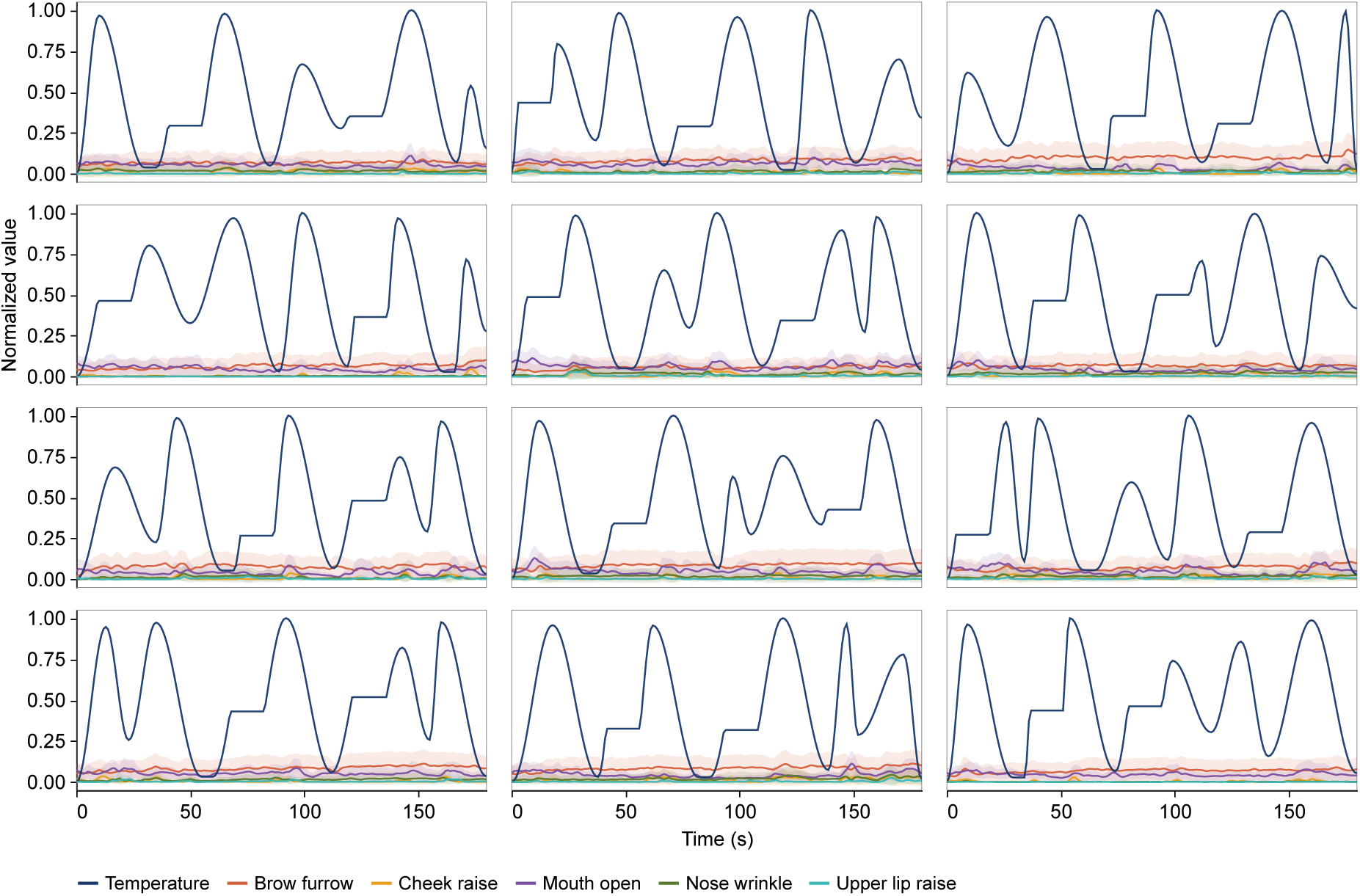
Grand-averaged means of facial expressions.

**Figure S6:**
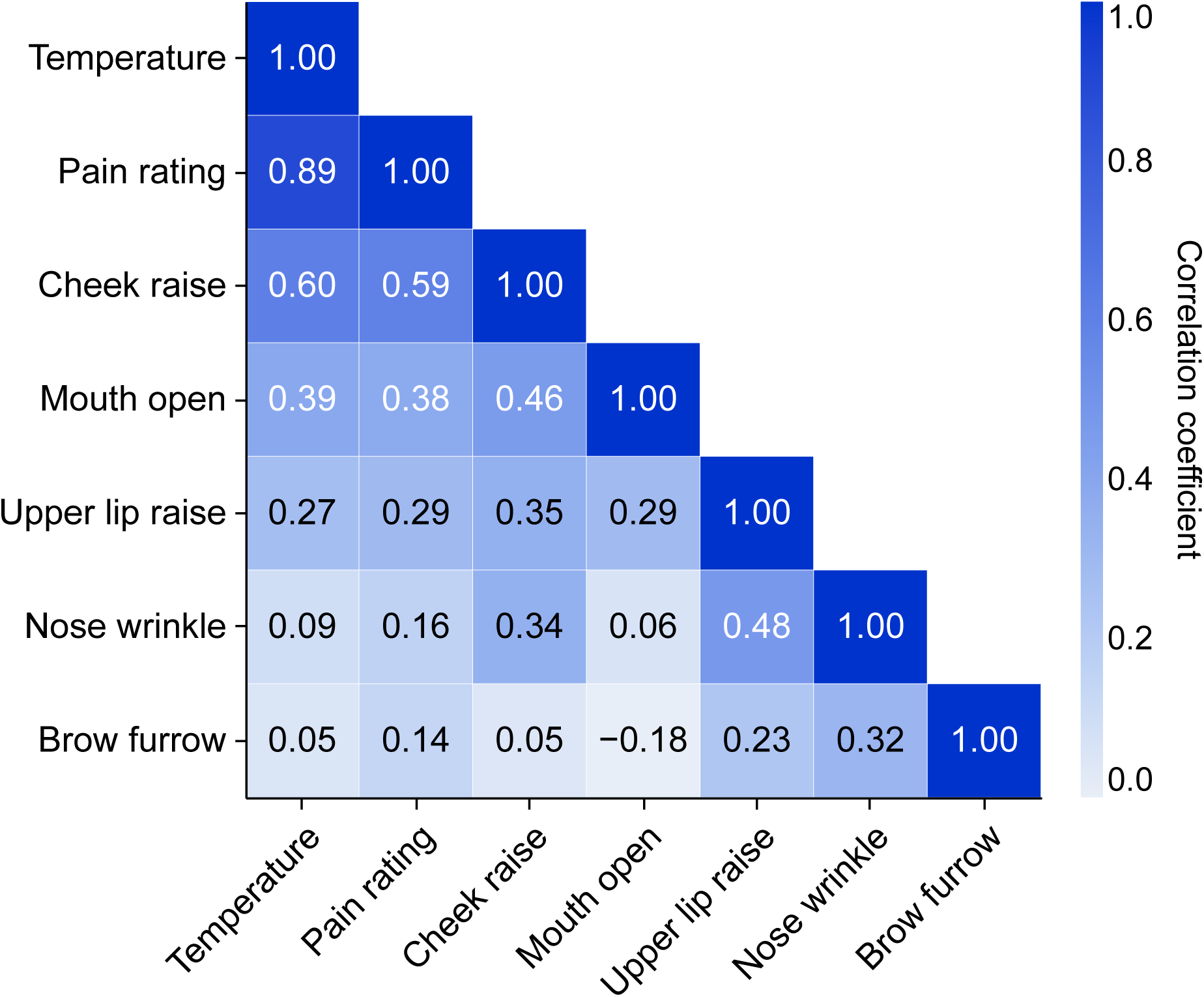
Correlation matrix of grand-averaged facial expressions, temperatures, and pain ratings. The heatmap shows Pearson correlation coefficients between the signals, with darker blue indicating stronger positive correlations.

**Table S5:** Temperature-referenced cross-correlation lags.

| Reference signal | Target signal | Mean lag (s) ± SD |
| --- | --- | --- |
| Temperature | Upper Lip Raise | 1.5 ± 7.5 |
| Temperature | Nose Wrinkle | 1.43 ± 4.41 |
| Temperature | Pupil Diameter | -0.07 ± 0.22 |
| Temperature | Mouth Open | -0.56 ± 0.71 |
| Temperature | Cheek Raise | -0.63 ± 0.87 |
| Temperature | EDA (tonic) | -0.95 ± 0.59 |
| Temperature | Pain Rating | -1.01 ± 0.34 |
| Temperature | Heart Rate | -1.76 ± 1.36 |
| Temperature | Brow Furrow | -2.45 ± 1.91 |
| Temperature | EDA (phasic) | -2.45 ± 0.89 |

*Note:* Positive values indicate the target signal precedes the temperature curve; negative values indicate the target signal follows the temperature curve. Values are presented as mean ± standard deviation across all grand-averaged stimulus functions.

**Table S6:**
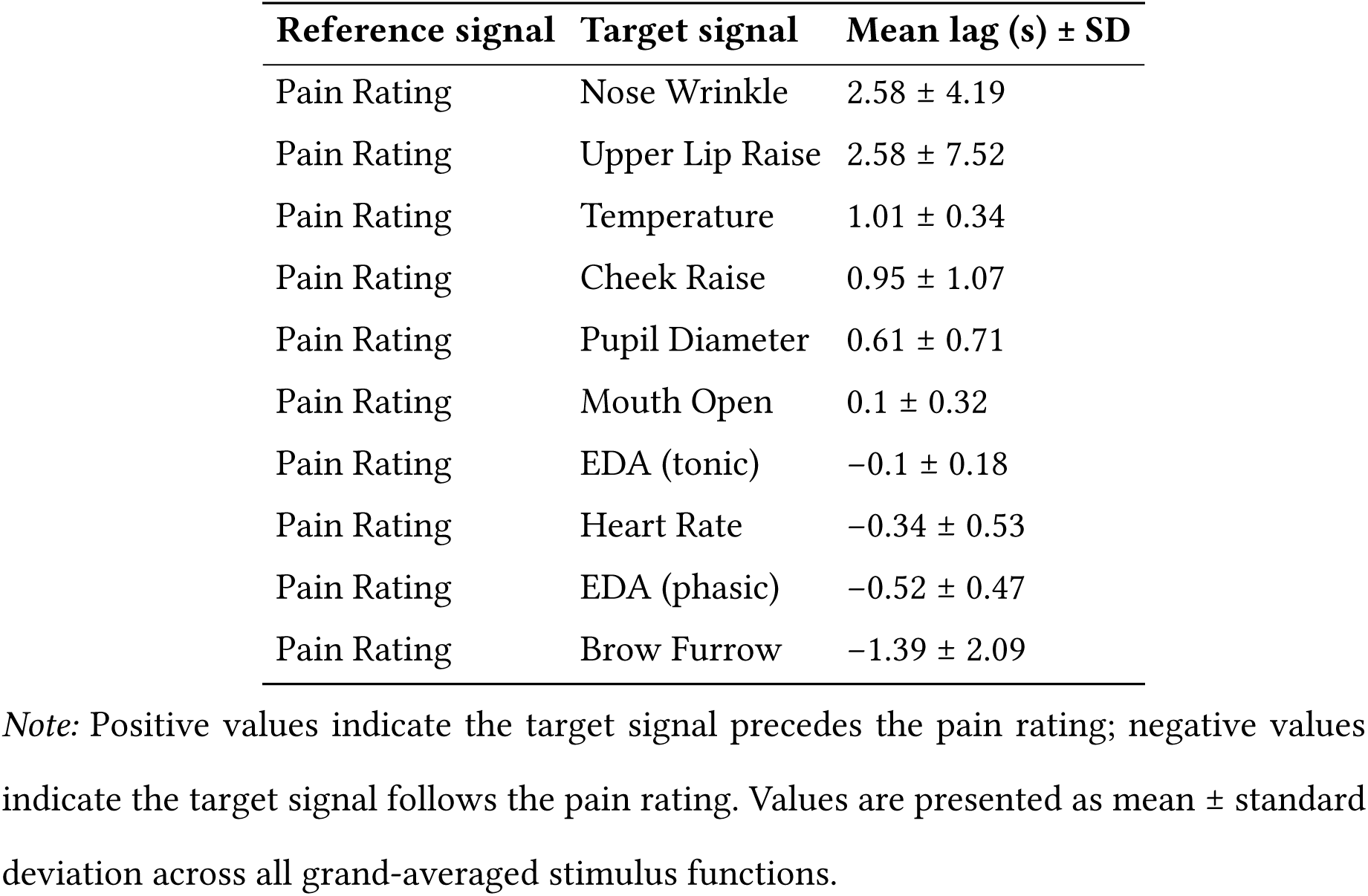
Pain rating-referenced cross-correlation lags.

| Reference signal | Target signal | Mean lag (s) ± SD |
| --- | --- | --- |
| Pain Rating | Nose Wrinkle | 2.58 ± 4.19 |
| Pain Rating | Upper Lip Raise | 2.58 ± 7.52 |
| Pain Rating | Temperature | 1.01 ± 0.34 |
| Pain Rating | Cheek Raise | 0.95 ± 1.07 |
| Pain Rating | Pupil Diameter | 0.61 ± 0.71 |
| Pain Rating | Mouth Open | 0.1 ± 0.32 |
| Pain Rating | EDA (tonic) | -0.1 ± 0.18 |
| Pain Rating | Heart Rate | -0.34 ± 0.53 |
| Pain Rating | EDA (phasic) | -0.52 ± 0.47 |
| Pain Rating | Brow Furrow | -1.39 ± 2.09 |

*Note:* Positive values indicate the target signal precedes the pain rating; negative values indicate the target signal follows the pain rating. Values are presented as mean ± standard deviation across all grand-averaged stimulus functions.

**Figure S7:**
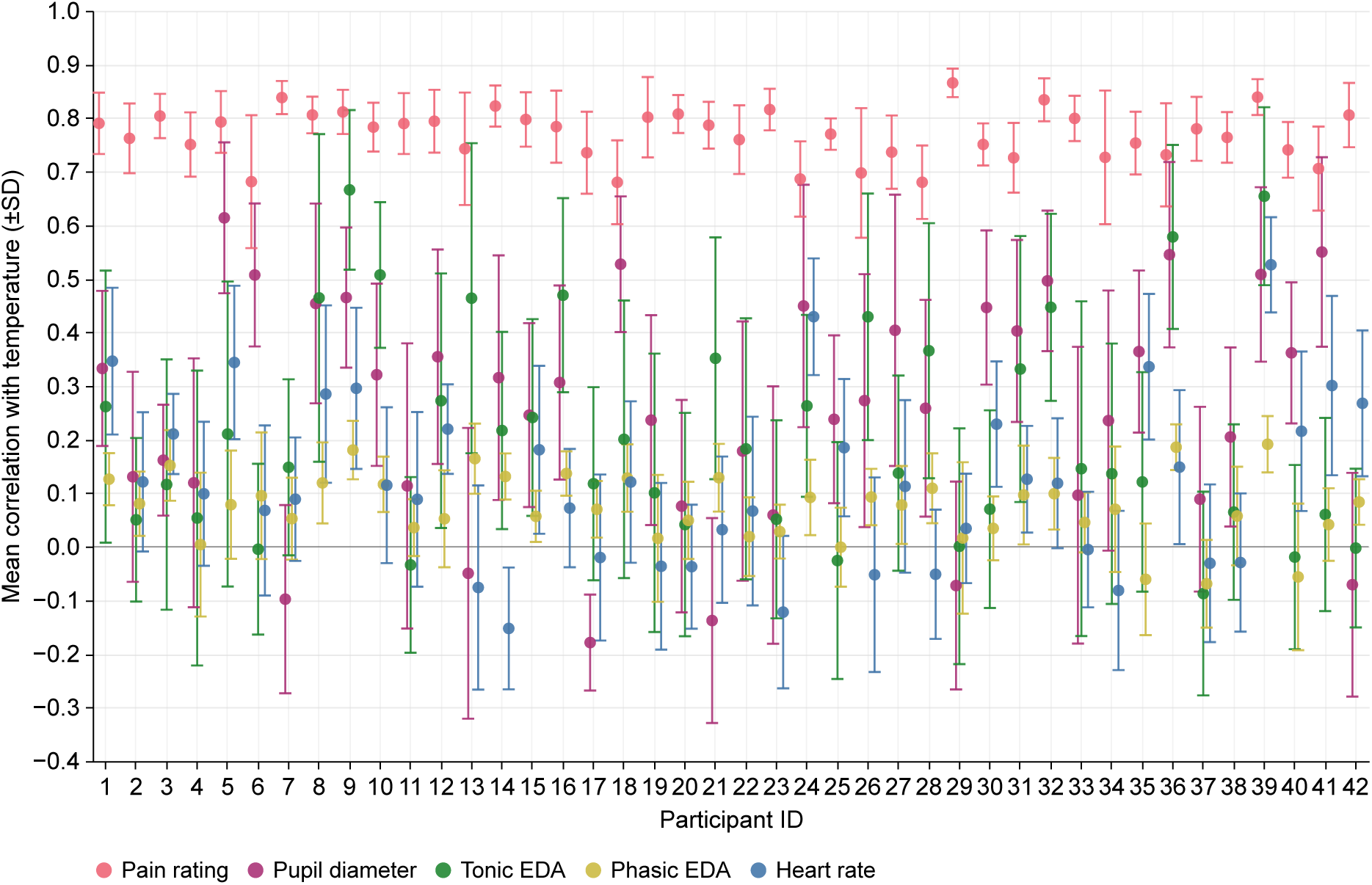
Correlations across participants between temperature and physiological signals.

**Figure S8:**
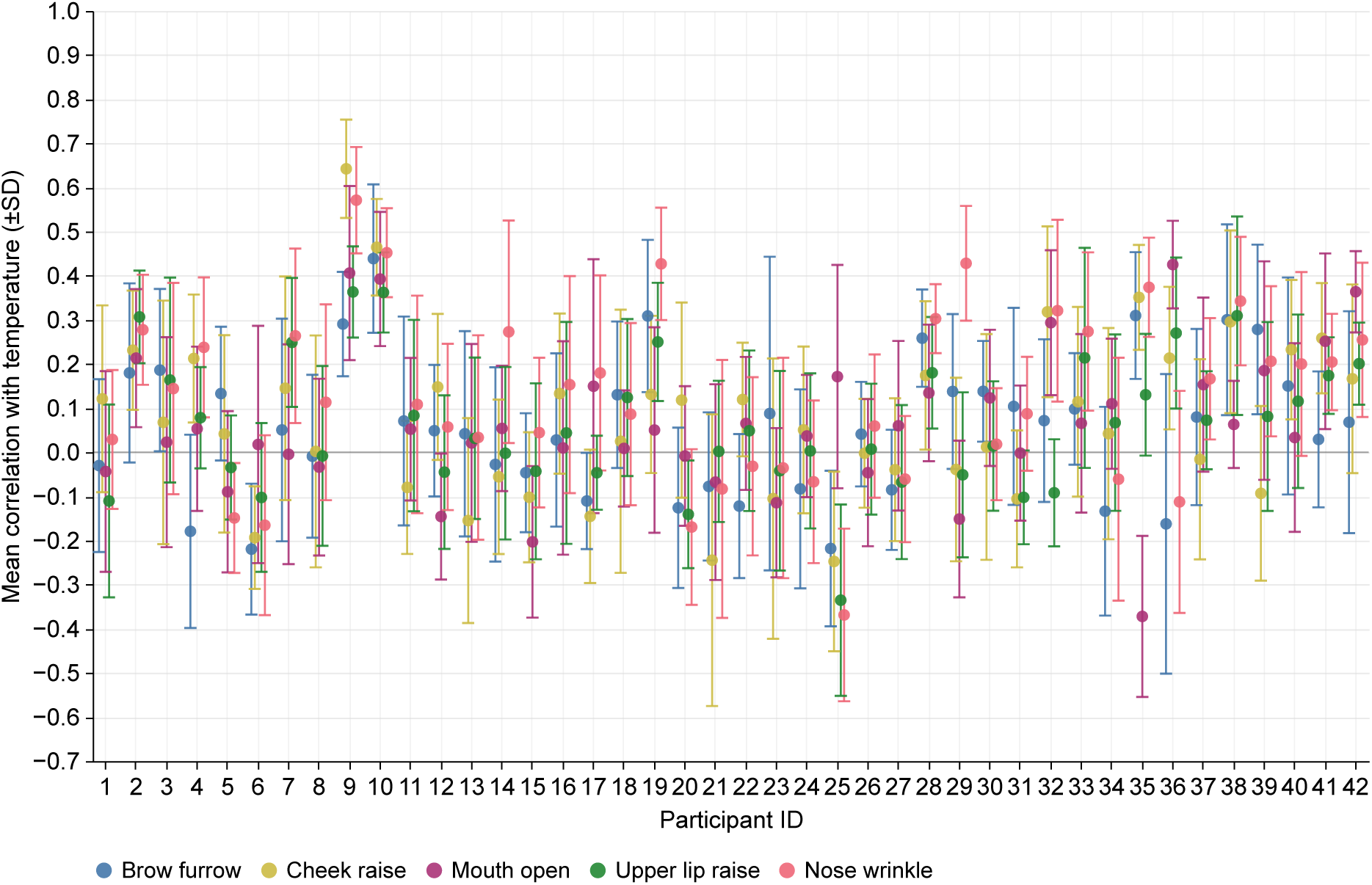
Correlations across participants between temperature and facial expressions.

#### Pain Ratings

**Figure S9:**
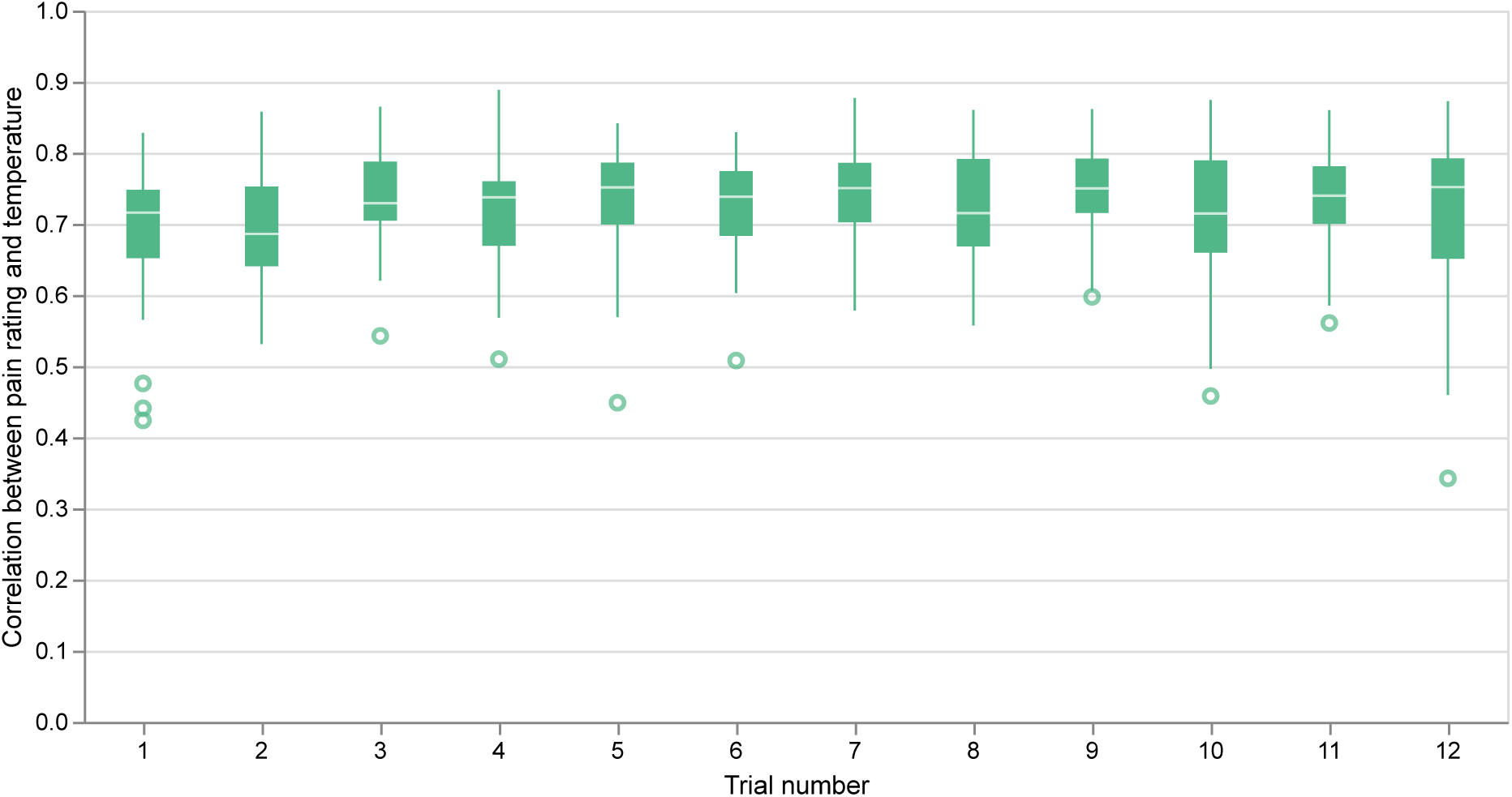
Correlation between pain ratings and temperature curves over the course of the experiment. Correlations between individual pain ratings and temperature curves remained stable across trials, with no meaningful learning effect observed.

**Figure S10:**
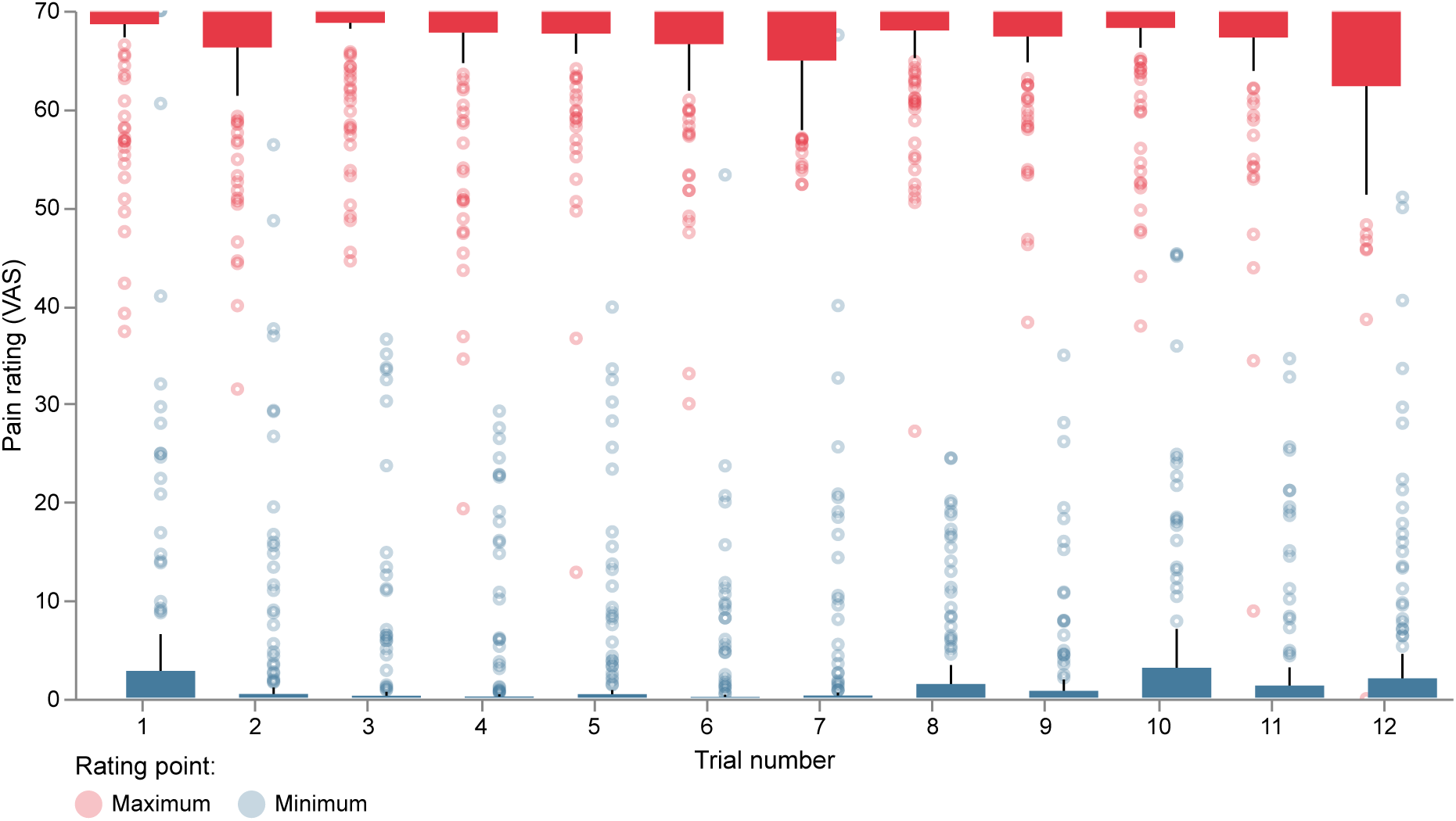
Maximum and minimum pain ratings in major decreasing intervals over the course of the experiment. With 1413 major decreasing temperature intervals across 471 trials, a total of 2826 rating points (one maximum and one minimum per decrease) are illustrated via the boxplots. Dots represent individual ratings; boxes represent the interquartile range with the median. Across all 12 trials, the median maximum pain rating was at the upper boundary of the scale (VAS 70), and the median minimum pain rating was at the lower boundary (VAS 0). This indicates that participants consistently identified and utilized the full extent of the rating scale, reflecting the expected pain peaks and troughs inherent in the stimulus design.

**Figure S11:**
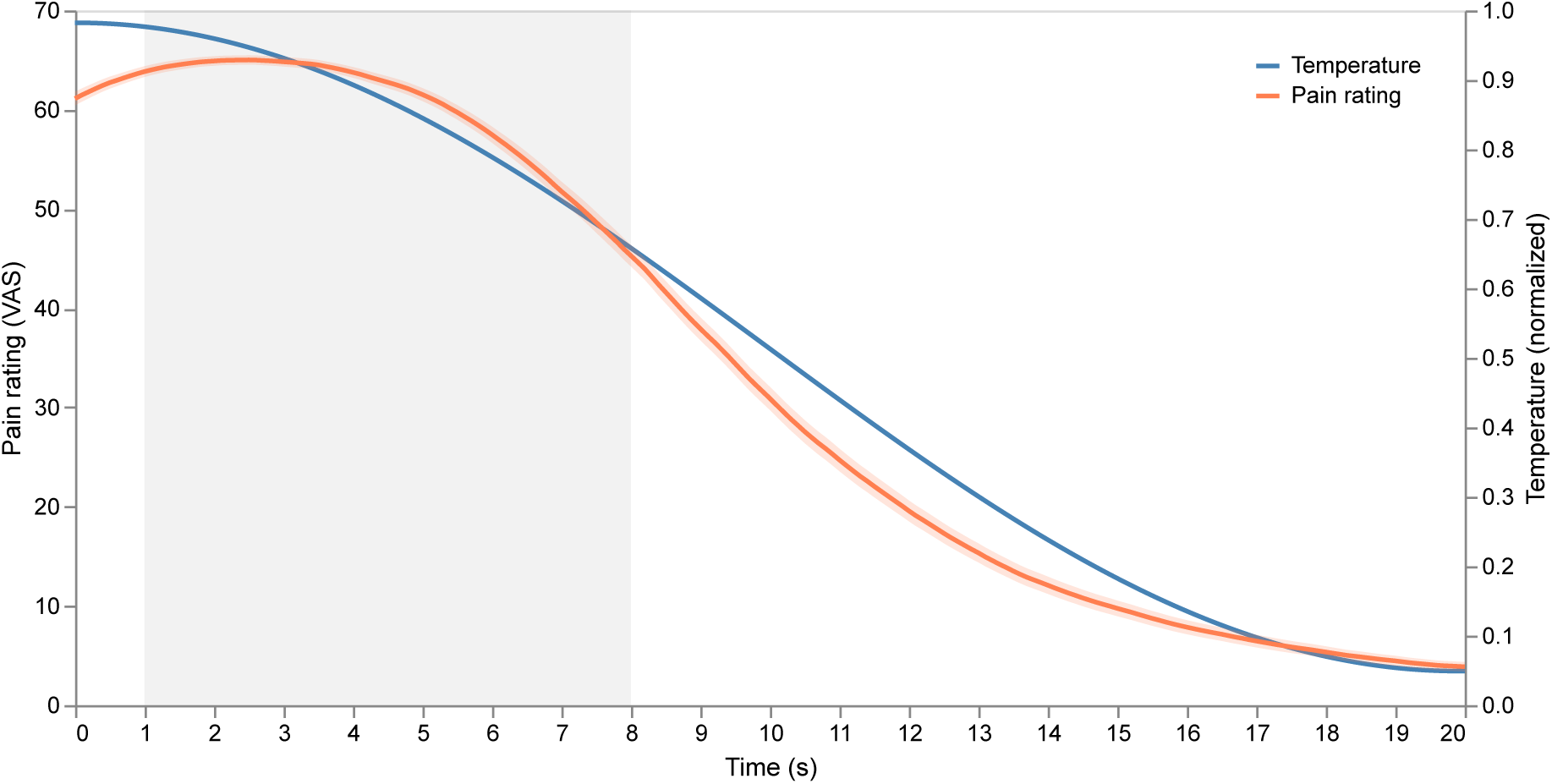
Average pain ratings over major temperature decreases. Pain ratings closely follow the temperature curve with a slight lag, reflecting psychophysical coupling. The grey shaded area (t = 1–8 s) shows the time window used for sample creation of decreases in the binary classification task. This window was chosen to capture the early phase of the decrease as timing is crucial for the proposed intervention rational.

### Supplementary Material F: Model Comparison and Optimization

#### Results

**Table S7:** Validation set accuracies across model architectures.

| Feature set | MLP | LightTS | TimesNet | PatchTST | iTransf. | Ensemble | EEGNet |
| --- | --- | --- | --- | --- | --- | --- | --- |
| EDA | 0.665 | 0.674 | 0.650 | **0.748** | 0.663 | - | - |
| HR | **0.609** | 0.596 | 0.572 | 0.575 | 0.585 | - | - |
| Pupil | 0.666 | 0.679 | 0.668 | 0.679 | **0.681** | - | - |
| EDA, HR | 0.655 | 0.676 | 0.617 | **0.756** | 0.628 | - | - |
| EDA, Pupil | 0.700 | 0.717 | 0.663 | **0.741** | 0.695 | - | - |
| EDA, HR, Pupil | 0.685 | 0.700 | 0.663 | **0.751** | 0.696 | - | - |
| Facial Expressions | 0.537 | 0.530 | 0.558 | **0.575** | 0.530 | - | - |
| Combined (w/o EEG) | 0.641 | 0.693 | 0.650 | **0.741** | 0.693 | 0.736 | - |
| EEG | 0.570 | 0.502 | - | - | **0.591** | - | 0.564 |
| EEG + EDA | 0.573 | 0.554 | - | - | 0.570 | **0.746** | - |
| Combined | 0.647 | 0.658 | - | - | 0.655 | **0.748** | - |

*Note:* Marked in bold are the highest validation accuracies. Ensemble stands for the ensemble model architecture of the specific feature set. iTransf. = iTransformer, HR = heart rate, EDA = electrodermal activity, EEG = electroencephalography. “Combined (w/o EEG)” comprises facial expressions, EDA, HR, and pupil diameter; “Combined” comprises those measures plus EEG.

**Table S8:**
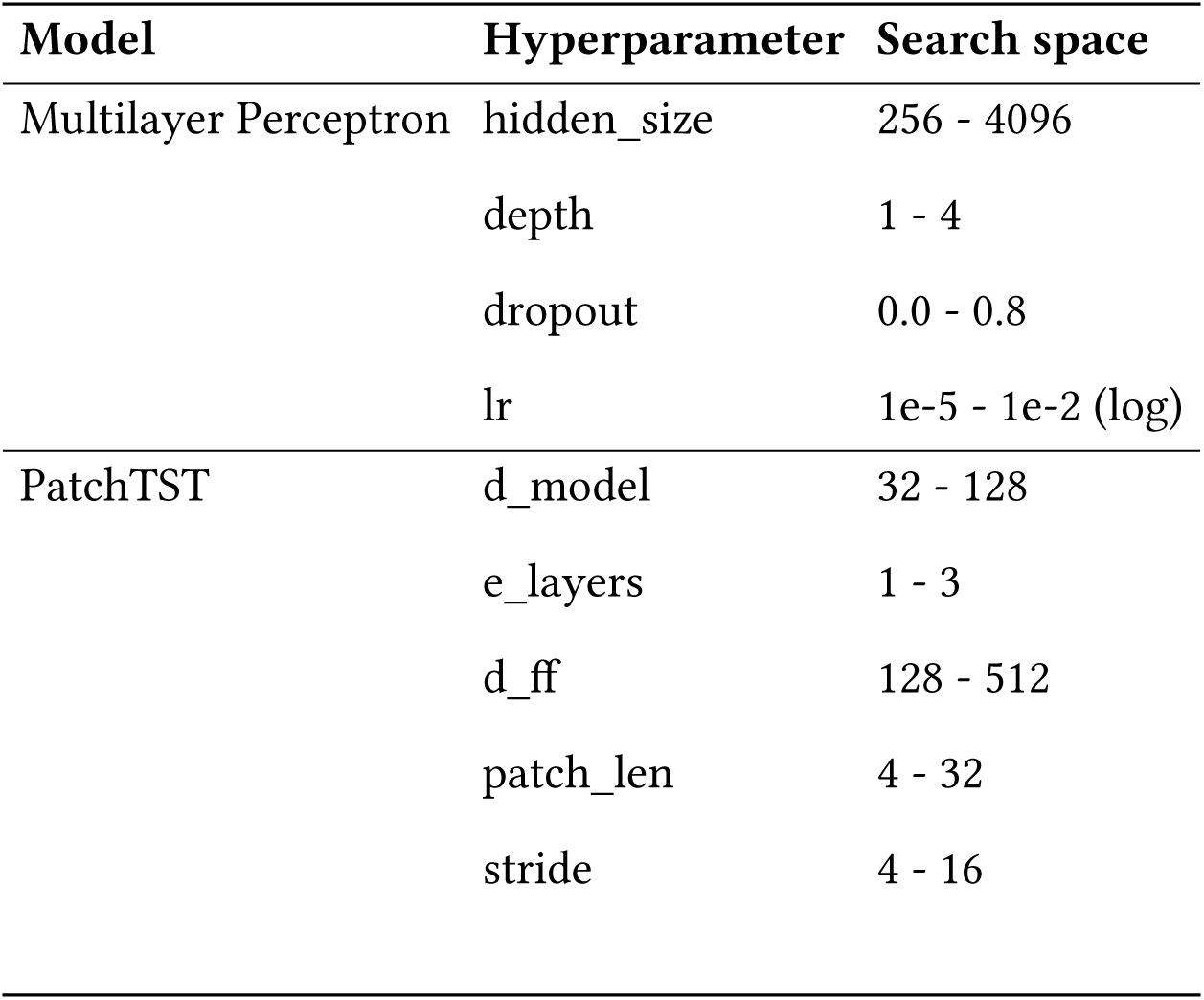
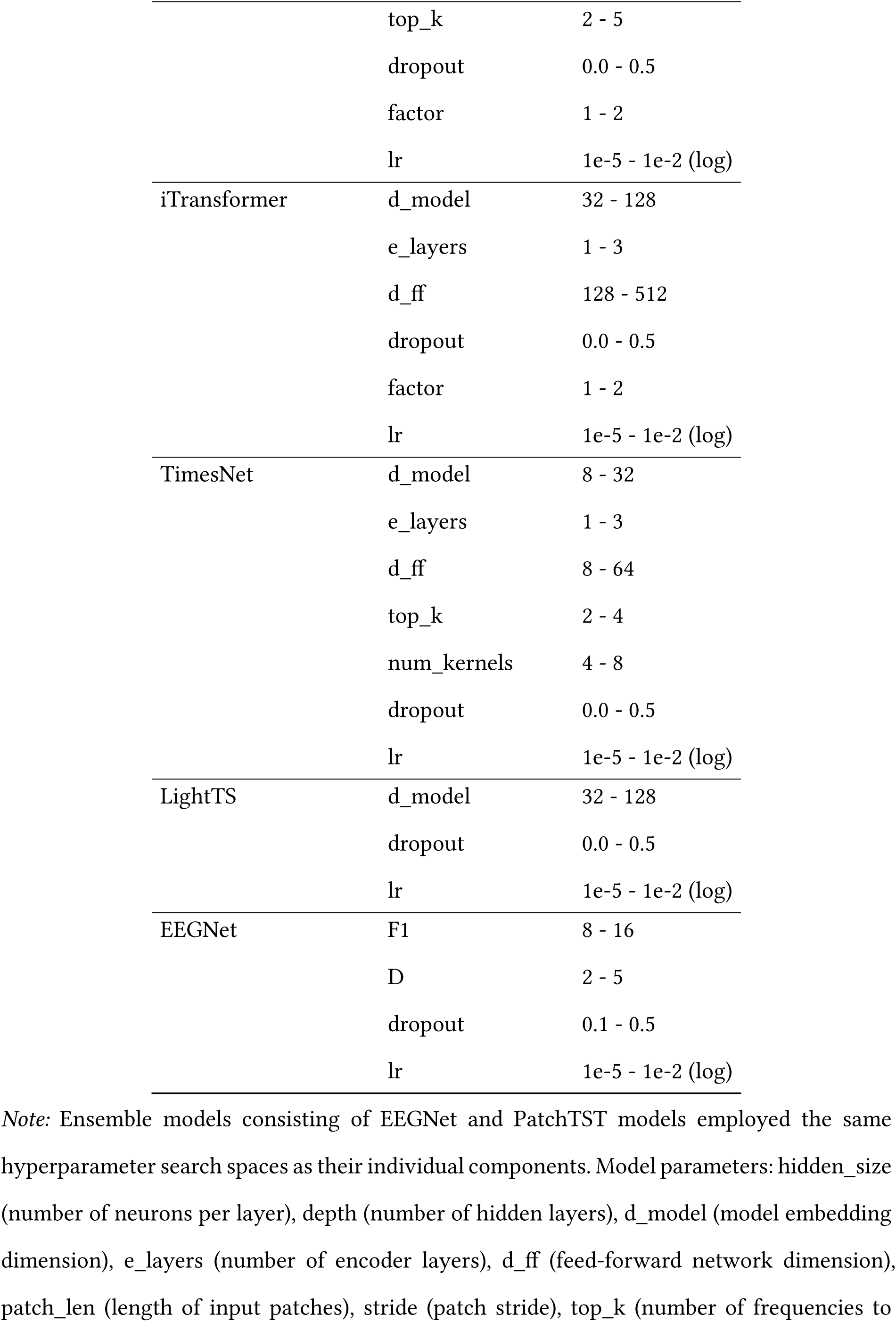
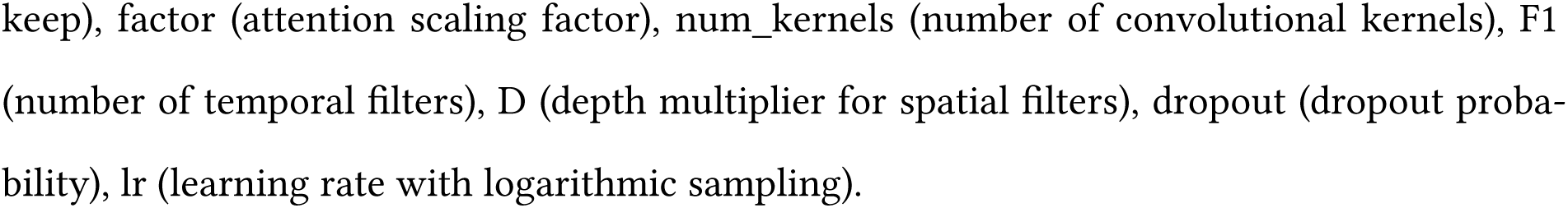
Hyperparameter search spaces.

**Table S9:**
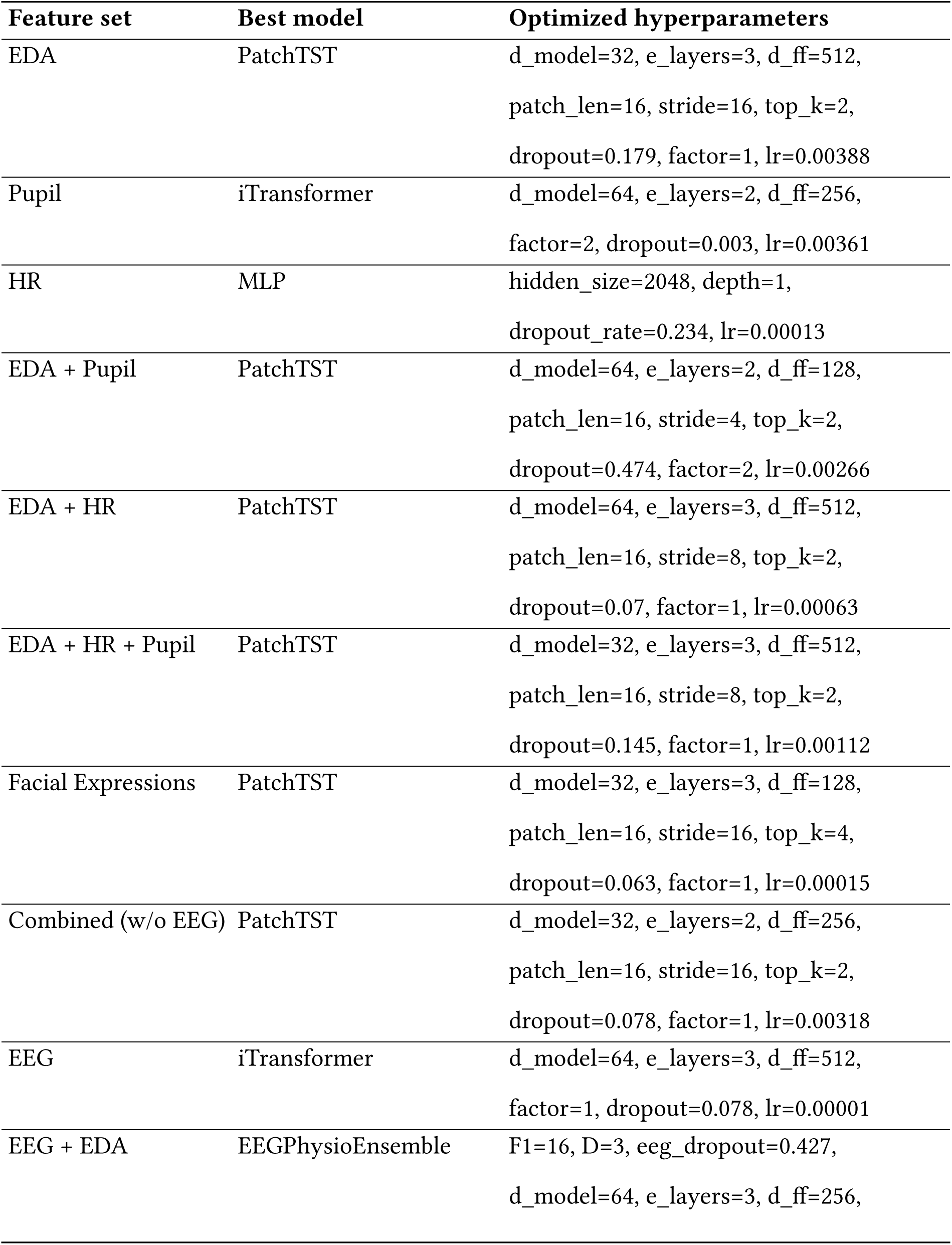
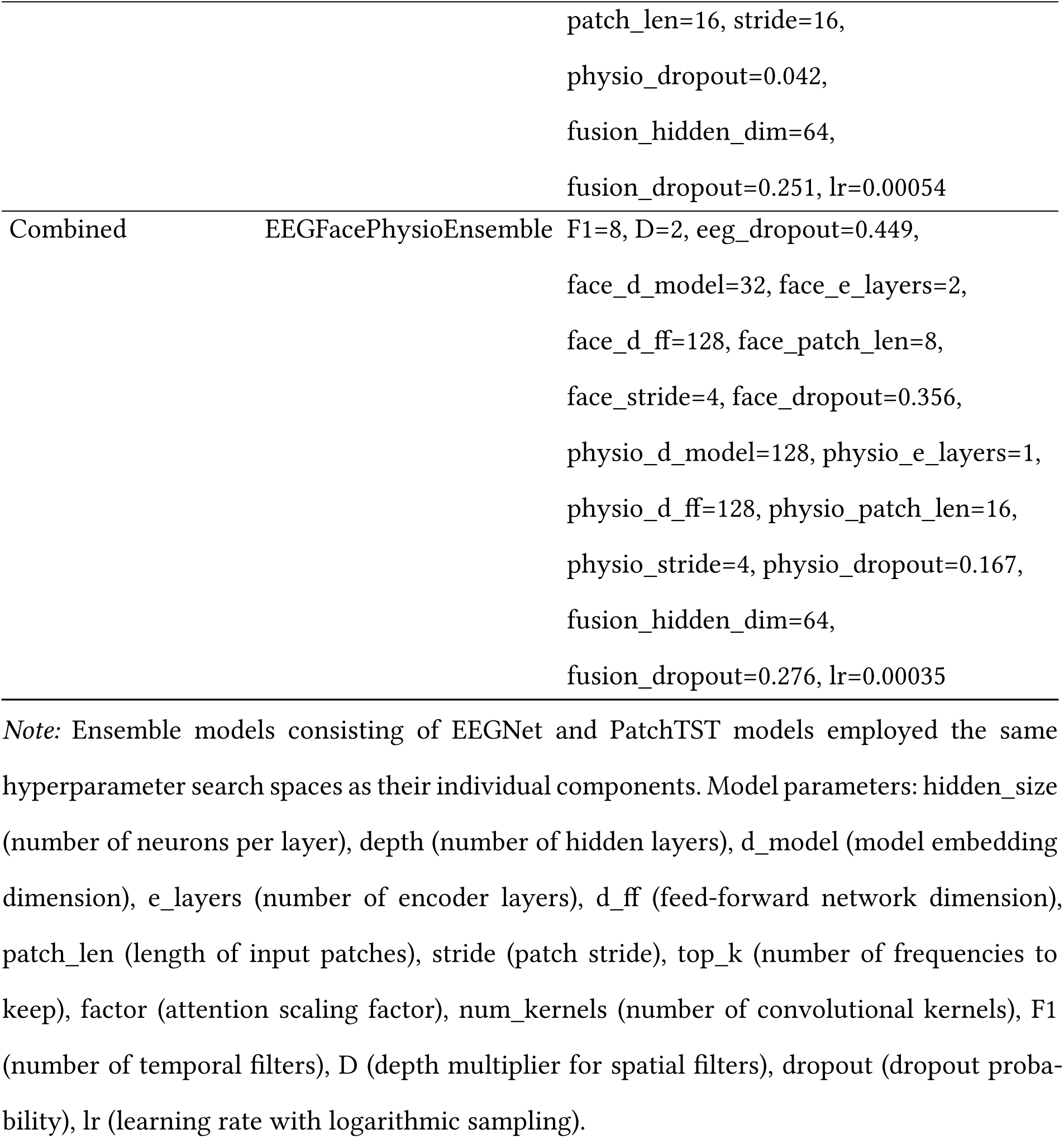
Optimized hyperparameters from the model comparison.

**Table S10:** Test set sample sizes for each participant.

| 1560 | Participant ID | Samples | Non-decreases | Decreases |
| --- | --- | --- | --- | --- |
| 1561 | 5 | 72 | 36 | 36 |
| 1562 | 9 | 78 | 42 | 36 |
| 1563 | 14 | 76 | 40 | 36 |
| 1564 | 20 | 69 | 33 | 36 |
| 1565 | 26 | 66 | 30 | 36 |
| 1566 | 27 | 64 | 28 | 36 |
| 1567 | 30 | 34 | 16 | 18 |
| 1568 | 31 | 51 | 24 | 27 |
| 1569 | 40 | 75 | 39 | 36 |
| 1570 | **total** | **585** | **288** | **297** |

**Table S11:** Test set accuracies box plot values.

| 1572 | Feature set | Mean | Median | Q1 | Q3 | IQR | Min | Max |
| --- | --- | --- | --- | --- | --- | --- | --- | --- |
| 1573 | EDA | 0.760 | 0.763 | 0.720 | 0.824 | 0.104 | 0.652 | 0.846 |
| 1574 | HR | 0.534 | 0.493 | 0.474 | 0.549 | 0.075 | 0.424 | 0.778 |
| 1575 | Pupil | 0.669 | 0.694 | 0.606 | 0.733 | 0.127 | 0.536 | 0.765 |
| 1576 | EDA + HR | 0.751 | 0.784 | 0.676 | 0.816 | 0.139 | 0.623 | 0.844 |
| 1577 | EDA + Pupil | 0.777 | 0.812 | 0.706 | 0.872 | 0.166 | 0.623 | 0.922 |
| 1578 | EDA + HR + Pupil | 0.764 | 0.766 | 0.676 | 0.846 | 0.170 | 0.636 | 0.902 |
| 1579 | Facial Expressions | 0.570 | 0.547 | 0.528 | 0.618 | 0.090 | 0.464 | 0.693 |
| 1580 | Combined (w/o EEG) | 0.745 | 0.797 | 0.706 | 0.806 | 0.100 | 0.609 | 0.824 |
| 1581 | EEG | 0.529 | 0.522 | 0.480 | 0.590 | 0.110 | 0.409 | 0.625 |
| 1582 | EEG + EDA | 0.734 | 0.750 | 0.706 | 0.792 | 0.086 | 0.560 | 0.824 |
| 1583 | Combined | 0.748 | 0.750 | 0.712 | 0.784 | 0.072 | 0.647 | 0.842 |

### Supplementary Material G: Model Inference Visualizations

**Figure S12:**
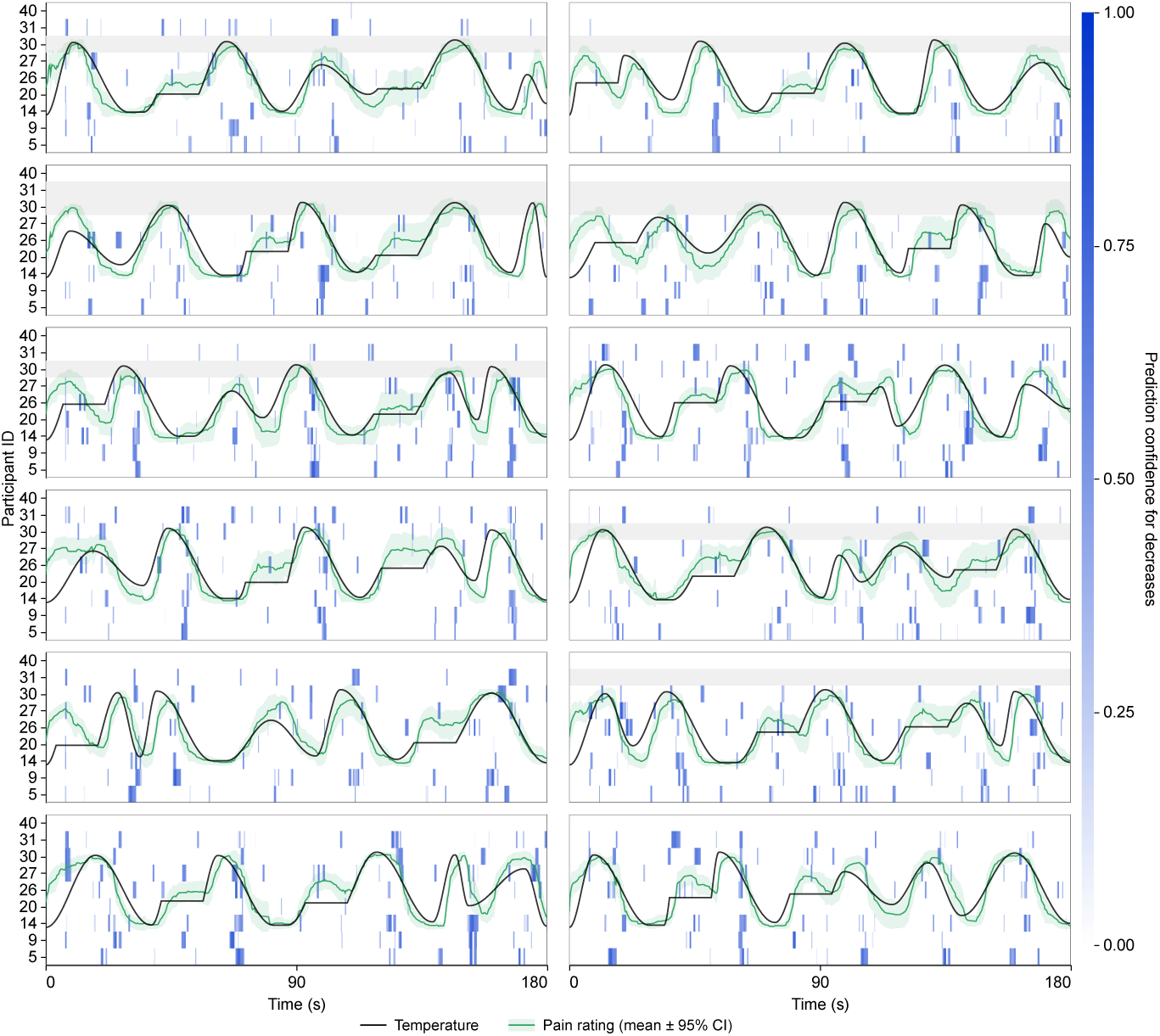
Continuous prediction heatmap for the decrease class using EDA. We applied the trained model using EDA, HR, and pupil diameter to all 12 continuous stimulus curves from the nine held-out participants. Each panel represents one stimulus curve; each heatmap row represents one participant. Blue intensity indicates the model’s predicted probability for the decrease class. Predictions at or above the threshold of 0.85 were classified as decreases. The black line shows the individually calibrated temperature curve; the green line and shaded band show the mean pain rating and 95% confidence interval across the test participants. Grey rows indicate missing trials. Pain ratings were recorded on a visual analogue scale from 0 to 70 and were not used during model training, which relied on temperature intervals as ground truth labels (decreases vs. increases and plateaus) and physiological signals as input. In Suppl. Figure S22 and Figure S23, we additionally provide prediction heatmaps for the detection of non-decreases and for both classes, respectively.

**Figure S13:**
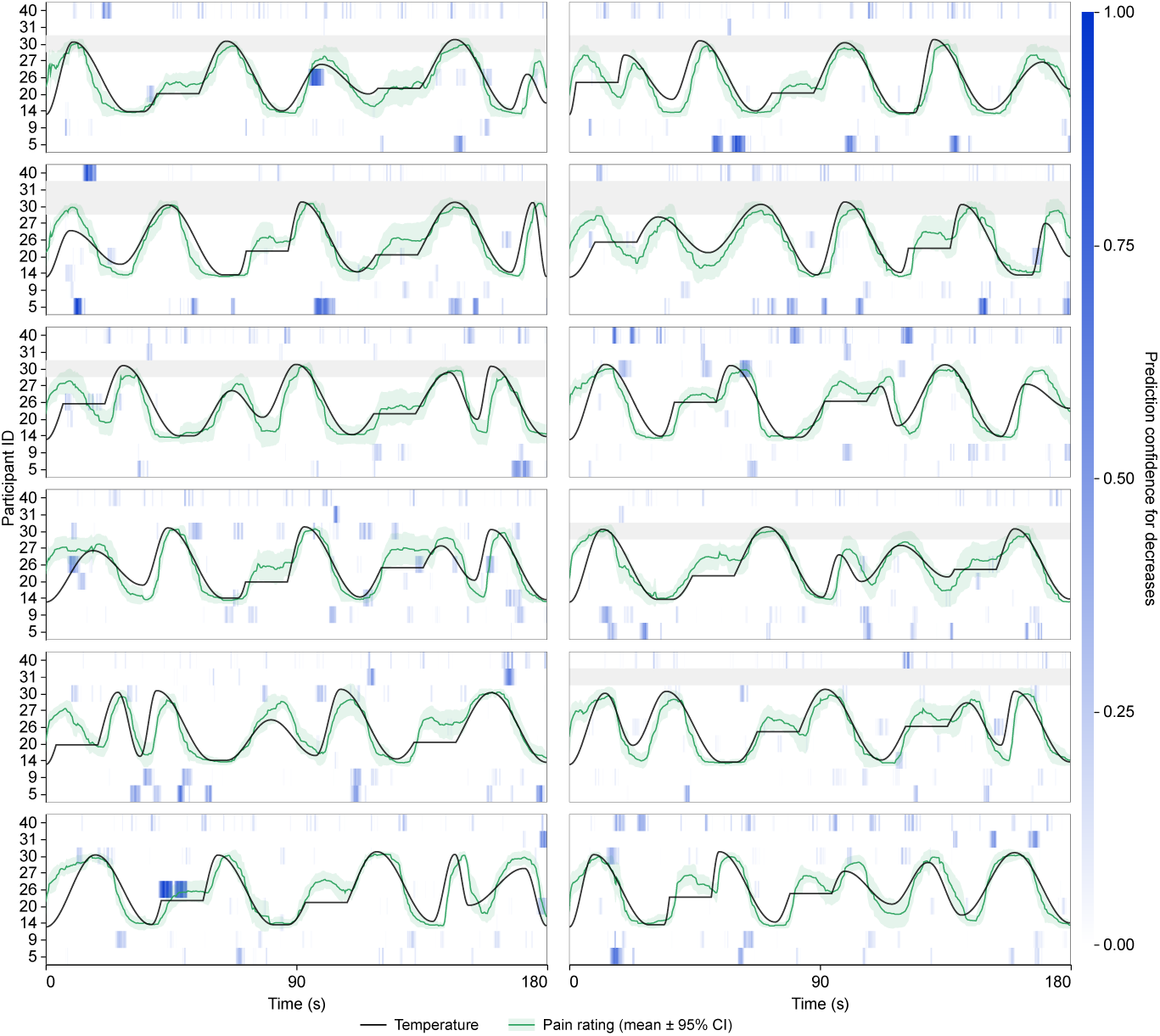
Continuous prediction heatmap for the decrease class using HR. Thresh-old = 0.60.

**Figure S14:**
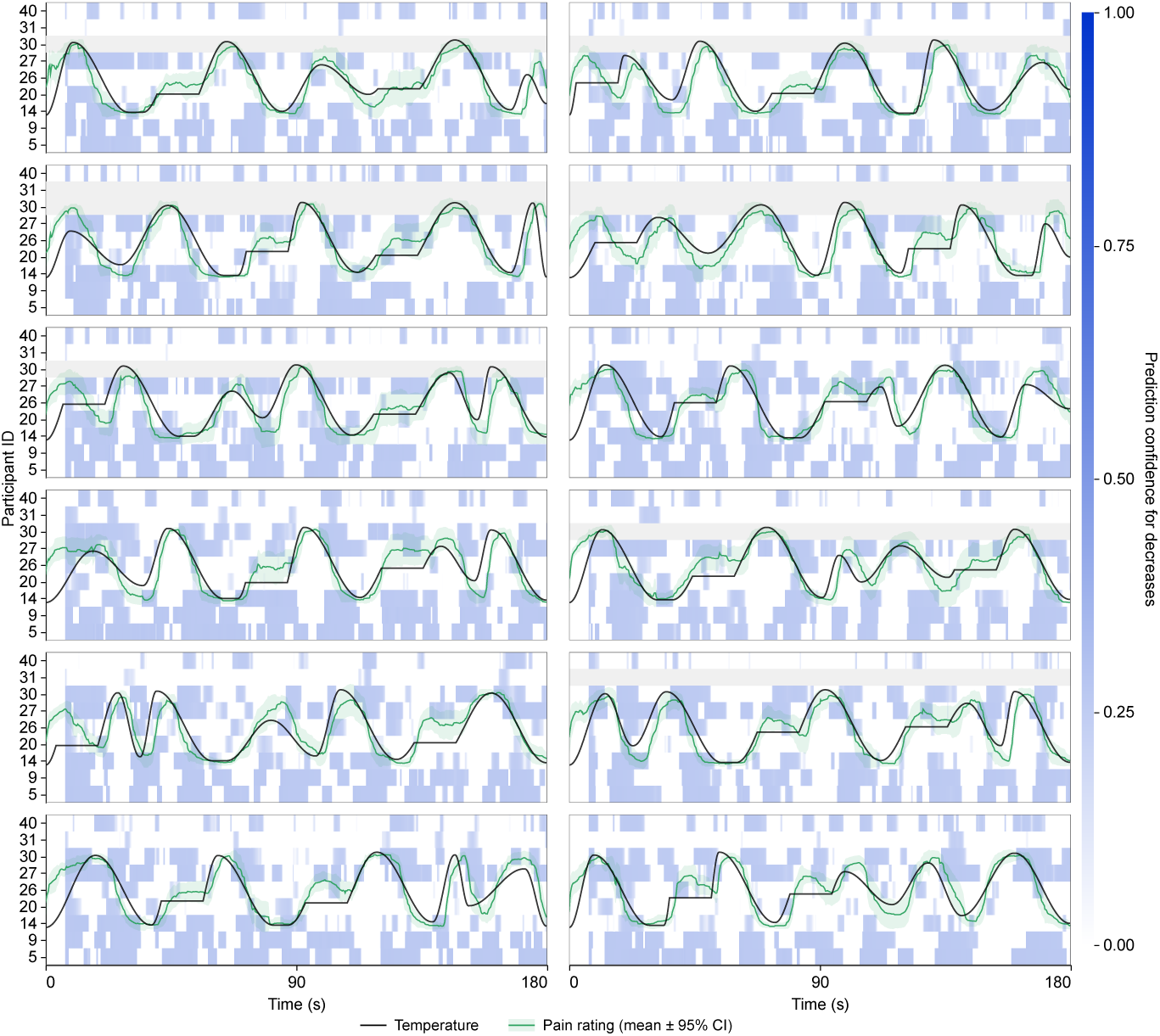
Continuous prediction heatmap for the decrease class using Pupil. Threshold = 0.50.

**Figure S15:**
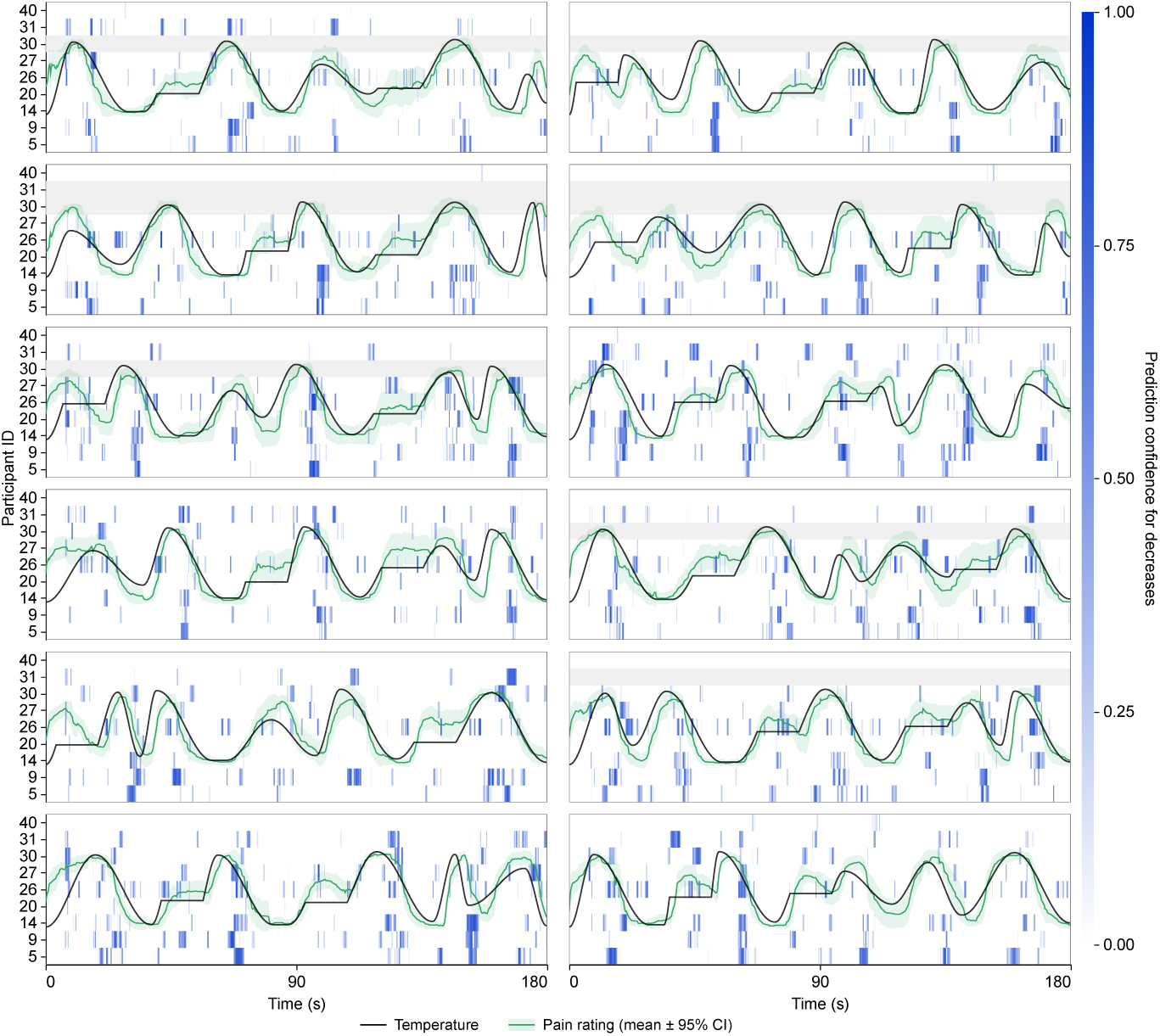
Continuous prediction heatmap for the decrease class using EDA + HR. Threshold = 0.85.

**Figure S16:**
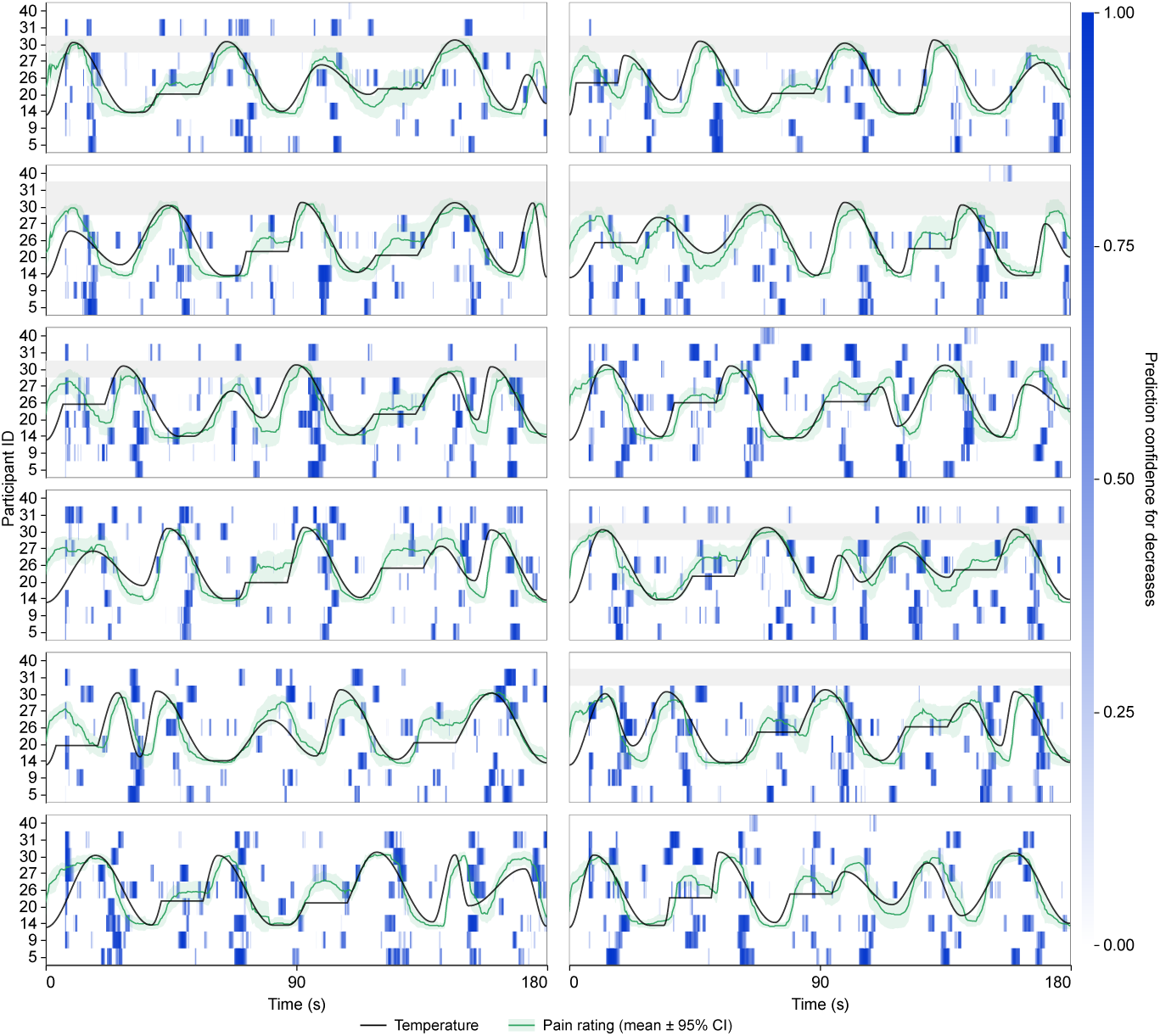
Continuous prediction heatmap for the decrease class using EDA + Pupil. Threshold = 0.90.

**Figure S17:**
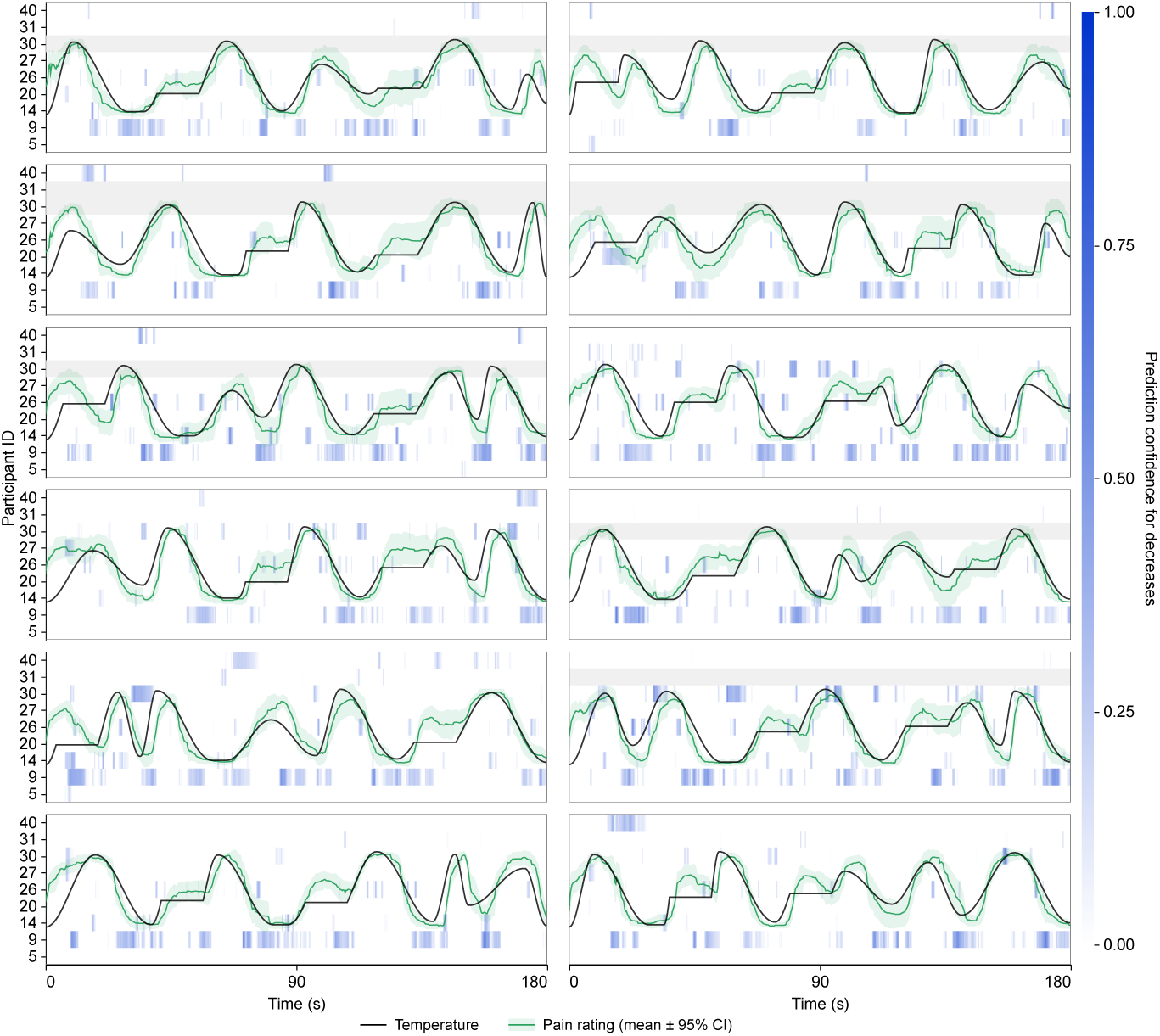
Continuous prediction heatmap for the decrease class using facial expres-sions. Threshold = 0.70.

**Figure S18:**
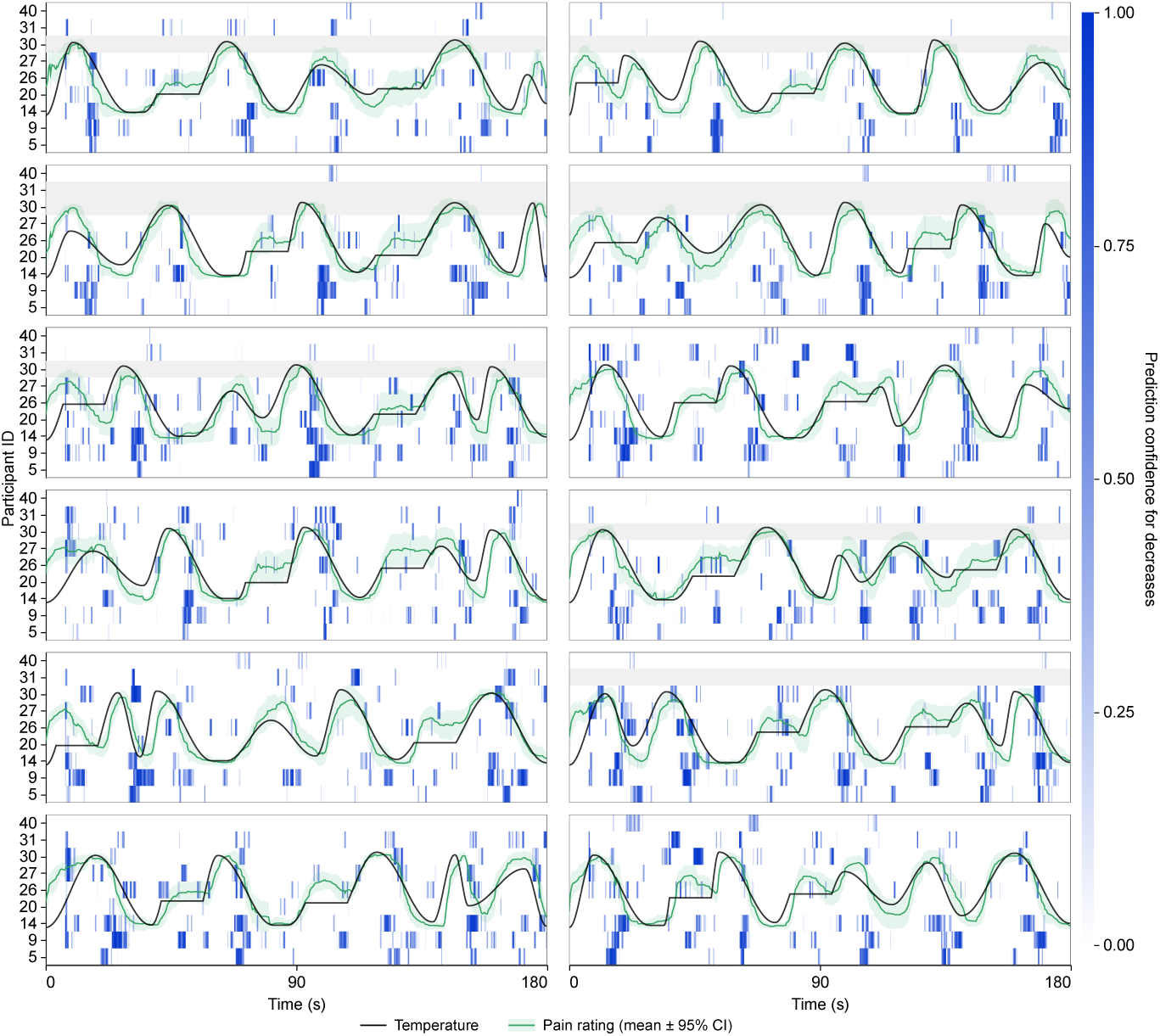
Continuous prediction heatmap for the decrease class using combined signals (without EEG). Threshold = 0.90. Includes facial expressions, EDA, HR, and Pupil.

**Figure S19:**
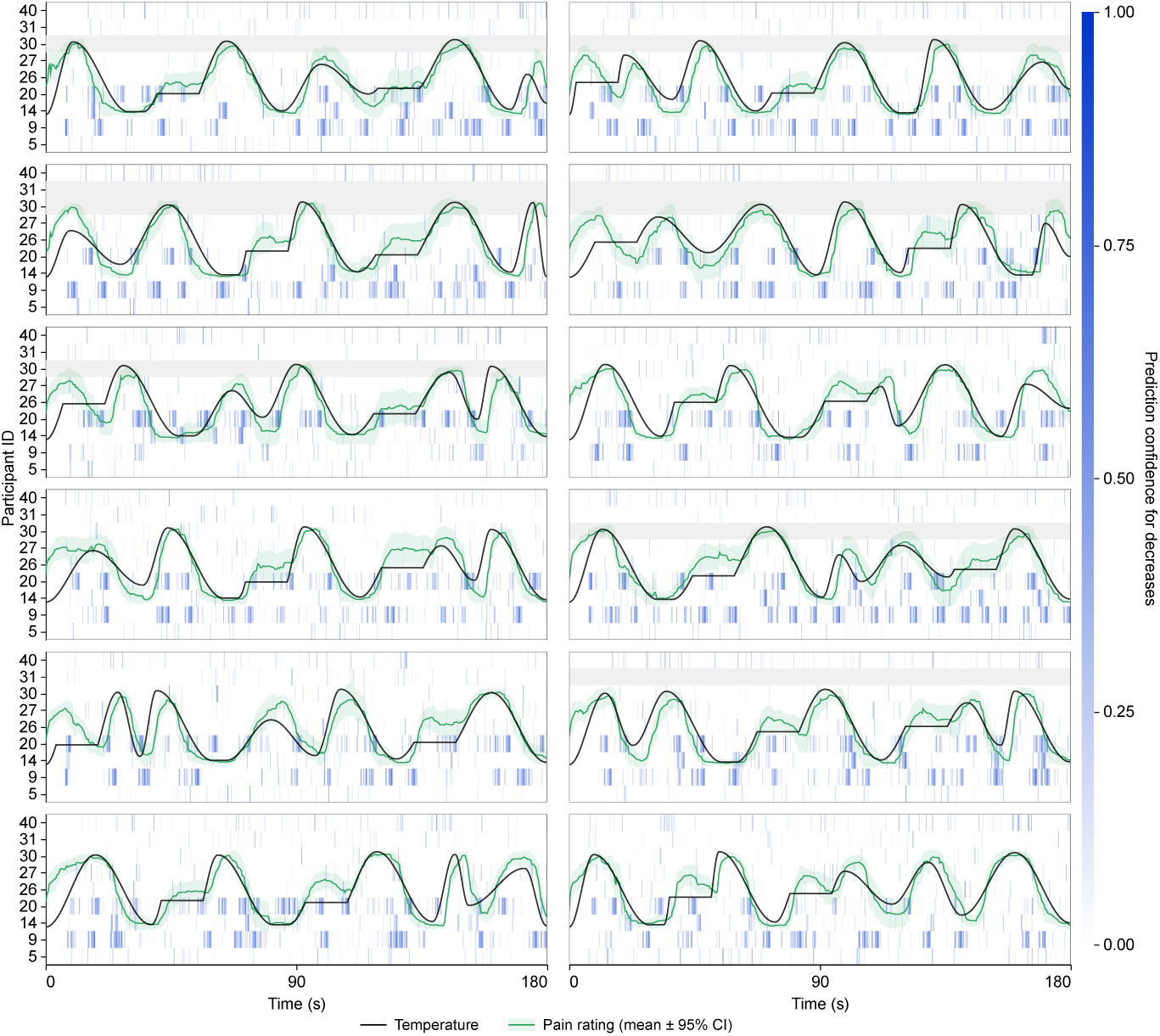
Continuous prediction heatmap for the decrease class using EEG. Thresh-old = 0.65.

**Figure S20:**
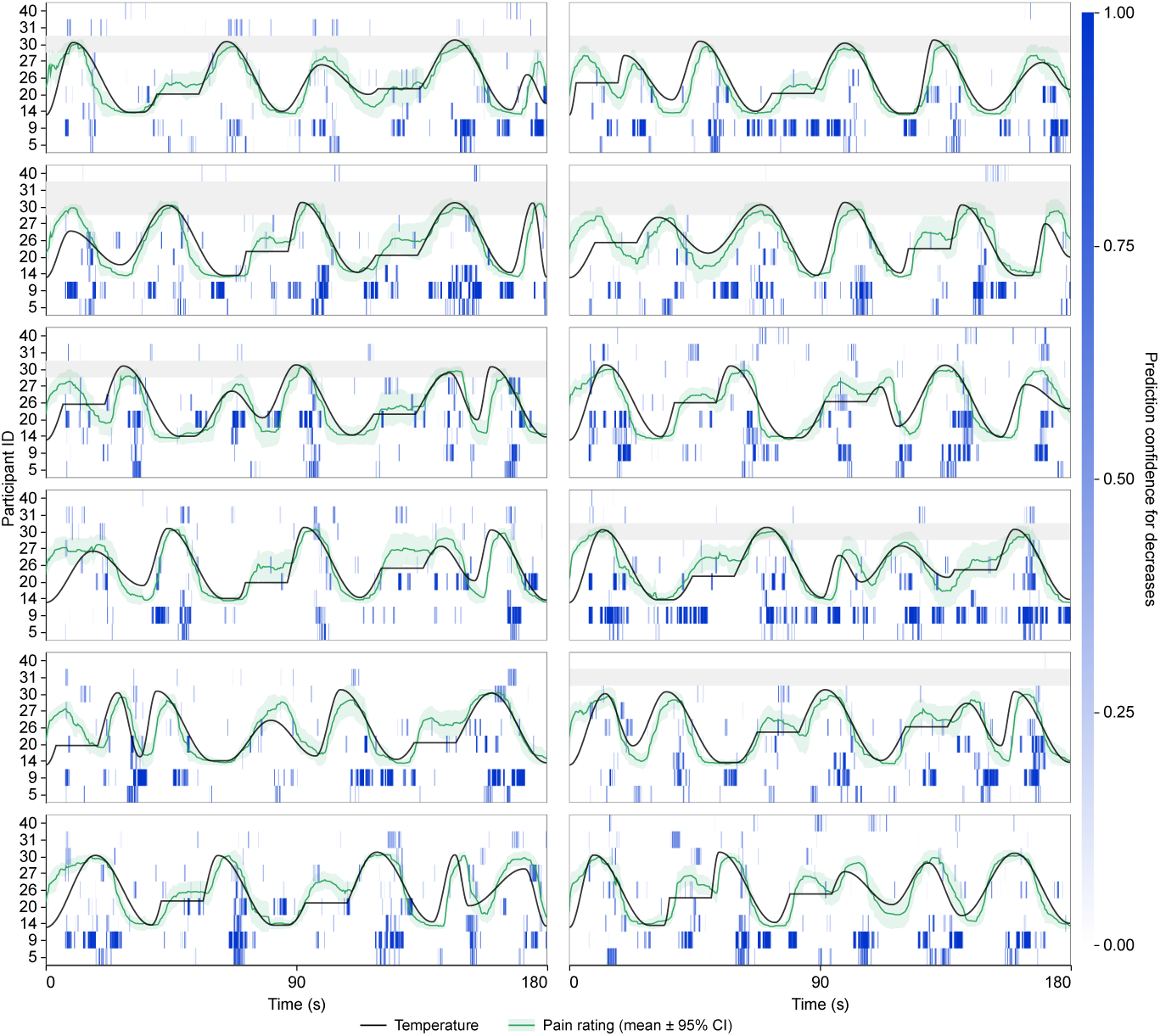
Continuous prediction heatmap for the decrease class using EEG + EDA. Threshold = 0.90.

**Figure S21:**
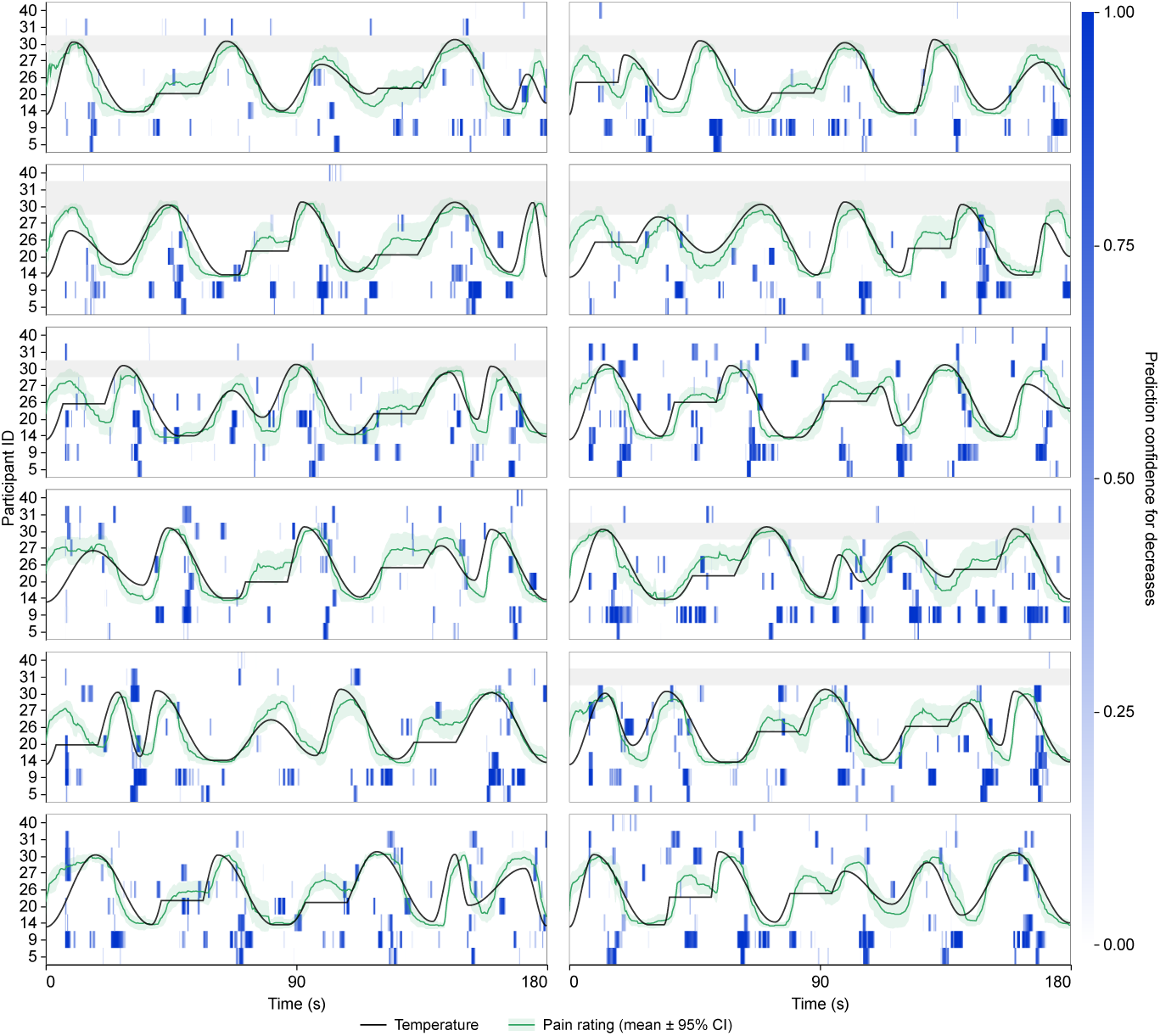
Continuous prediction heatmap for the decrease class using combined signals. Threshold = 0.90. Includes EEG, facial expressions, EDA, HR, and pupil diameter.

**Figure S22:**
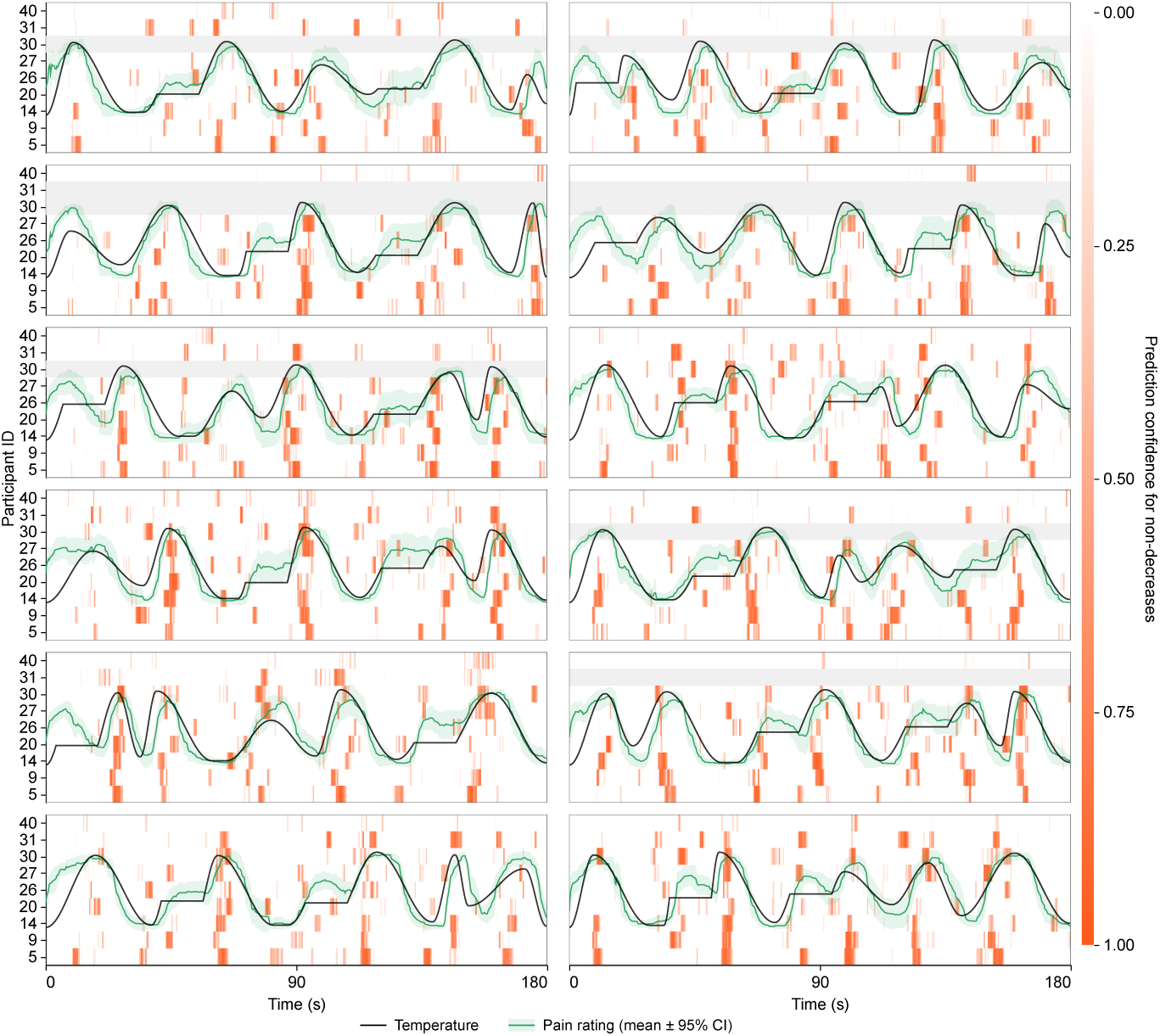
Continuous prediction heatmap for the non-decrease class using EDA + HR + Pupil. Threshold = 0.90.

**Figure S23:**
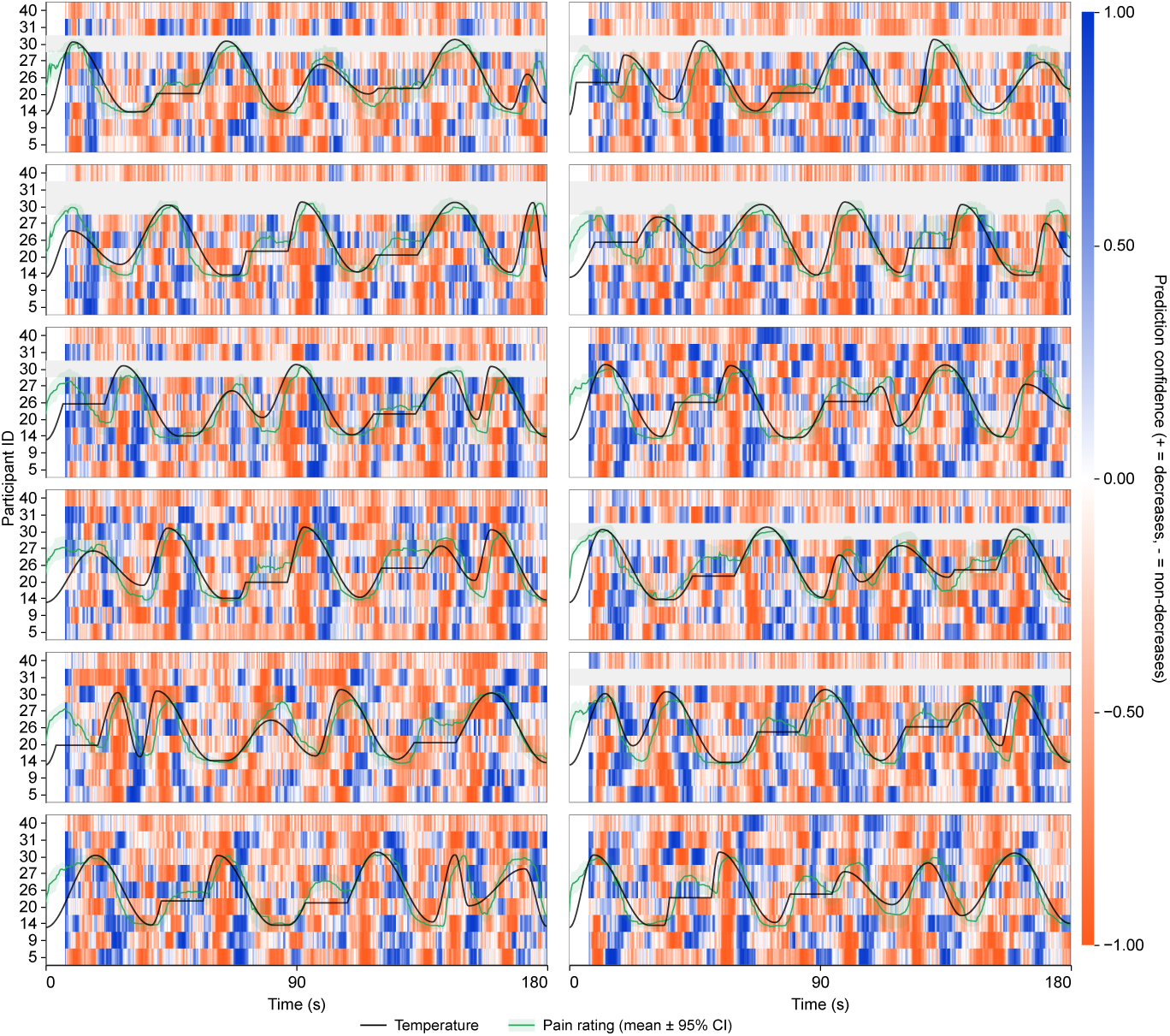
Continuous prediction heatmap for the decrease– and non-decrease classes using EDA + HR + Pupil. Threshold = 0.50.

